# The fungal ligand chitin directly binds and signals inflammation dependent on oligomer size and TLR2

**DOI:** 10.1101/270405

**Authors:** Katharina Fuchs, Yamel Cardona Gloria, Olaf-Oliver Wolz, Franziska Herster, Lokesh Sharma, Carly Dillen, Christoph Täumer, Sabine Dickhöfer, Zsofia Bittner, Truong-Minh Dang, Anurag Singh, Daniel Haischer, Maria A. Schlöffel, Kirsten J. Koymans, Tharmila Sanmuganantham, Milena Krach, Nadine A. Schilling, Felix Frauhammer, Lloyd Miller, Thorsten Nürnberger, Salomé LeibundGut-Landmann, Andrea A. Gust, Boris Macek, Martin Frank, Cécile Gouttefangeas, Charles S. Dela Cruz, Dominik Hartl, Alexander N.R. Weber

**Author notes:** **Contact information (Corresponding Author and Lead Contact)** Alexander N. R. Weber, Interfaculty Institute for Cell Biology, Department of Immunology, University of Tübingen, Auf der Morgenstelle 15, 72076 Tübingen, Germany. contributed equally (Katharina Fuchs, Yamel Cardona Gloria).

## Abstract

Chitin is a highly abundant polysaccharide and linked to fungal infection and asthma. Unfortunately, its polymeric structure has hampered the identification of immune receptors directly binding chitin and signaling immune activation and inflammation, because purity, molecular structure and molarity are not well definable for a polymer typically extracted from biomass. Therefore, by using defined chitin (N-acetyl-glucosamine) oligomers, we identified six subunit long chitin chains as the smallest immunologically active motif and the innate immune receptor Toll-like receptor (TLR) 2 as the primary fungal chitin receptor on human and murine immune cells. Chitin oligomers directly bound TLR2 with nanomolar affinity and showed both overlapping and distinct signaling outcomes compared to known mycobacterial TLR2 ligands. Conversely, chitin oligomers shorter than 6 subunits were inactive or showed antagonistic effects on chitin/TLR2-mediated signaling, hinting to a size-dependent sensing/activation system unexpectedly conserved in plants and humans. Since blocking the chitin-TLR2 interaction effectively prevented chitin-mediated inflammation *in vitro* and *in vivo*, our study highlights the chitin TLR2 interaction as a potential target for developing novel therapies in chitin-related pathologies and fungal disease.

## Introduction

Fungal infections and airway inflammation in allergic asthma are associated with an immense socio-economic burden globally (Brown et al, 2012; To et al, 2012) and have both been linked to immune activation by the second most abundant polysaccharide in nature, chitin. Chitin is a β-(1-4)-N-acetyl D-glucosamine (NAG, also referred to as GlcNAc) homo-polymer (Morgulis, 1916) and is found in fungi, parasitic nematodes, the crustaceans and insects, and house dust mite allergen (Lee et al, 2008; Mack et al, 2015). Chitin is known to elicit strong immunogenic activity with particular relevance for fungal infection (Bueter et al, 2013; Cunha et al, 2010) and airway inflammation during asthma (Choi et al, 2016; Van Dyken et al, 2017), and constitutes a highly conserved microbe-associated molecular pattern (MAMP) (Lee et al, 2008) eliciting immune activation in plants and mammals. In the latter chitin is considered an ‘orphan MAMP’ by many researchers in the field as its direct immune receptor has so far not been identified. MAMPs are typically sensed by direct binding to pattern recognition receptors (PRRs) and in mammals the best-studied PRR families include Toll-like receptors (TLRs) and C-type-lectin-like receptors (CLRs) (Hoving et al, 2014; Kawasaki & Kawai, 2014). Certain CLRs detect oligosaccharide structures common on fungal pathogens such as β-glucans which are sensed by the CLR Dectin-1 (Hoving et al, 2014); TLRs directly recognize non-oligosaccharide and more structurally diverse MAMPs – e.g. lipopeptides (via TLR2), lipopolysaccharide (LPS, via TLR4) or nucleic acids (via TLRs 7, 8 and 9) – via their extracellular domains (ECD) and initiate cytoplasmic signaling involving the adapter myeloid differentiation factor 88 (MyD88). Whereas respiratory burst and phagocytosis are triggered only by CLRs, CLR and TLR signaling may cooperate for pro-inflammatory cytokine and/or interferon (IFN) transcription and the initiation of adaptive immunity (Kawasaki & Kawai, 2014).

Although plants lack an adaptive immune response, they also succumb to fungal infections and feature a sophisticated innate immune system which shows interesting similarities to mammalian innate immunity with regard to MAMP recognition, for example, regarding flagellin and chitin (Jones & Dangl, 2006; Kawasaki & Kawai, 2014). Fungal chitin was shown to act via the composite CEBiP/CERK1 chitin receptor system in both rice (*Oryza sativa*) and *Arabidopsis thaliana*. Interestingly, this required a minimum chitin oligomer length for target gene (e.g. *FRK1)* induction and antimicrobial ROS production (Hayafune et al, 2014; Liu et al, 2012), a feature not reported for mammalian chitin recognition thus far: although in mammals immunological effects have been considered size-dependent such size-dependence was only studied on a macromolecular and not oligomer or molecular, scale (Alvarez, 2014; Da Silva et al, 2009). For example, even the smallest (however, typically ∼1 μm) macromolecular chitin to be considered immuno-stimulatory to date can be estimated to contain hundreds to thousands of NAG subunits, thus exceeding the size of entire human immune cells. Chitin <0.2 μm has been considered immunologically inactive in the field and thus irrelevant in mammals (Lee et al, 2008). Since oligomeric chitin would better match the dimensions of the PRRs discussed so far, this conclusion seems counterintuitive. However, it has not been properly challenged due to the difficulty in obtaining well-defined oligomeric chitin, whereas crude chitin of poorly defined purity, molecular structure and/or molarity is readily available from fungal or shrimp biomass. Studies of such crude chitin in PRR knockout (KO) mice have thus yielded a controversial list of putative chitin receptors in mammals, but direct binding of chitin to any of these receptors has not been demonstrated so far. The receptor(s) for the observed pleiotropic immunological effects of chitin in mammalian immune cells is therefore considered elusive (Bueter et al, 2013). More than a century after the initial report of its chemical composition (Morgulis, 1916), critical properties of chitin immune recognition – a binding receptor and its molecular size-preference – in mammals thus still remain elusive.

Using defined chitin oligomers rather than crude chitin we here show TLR2 to be a mammalian immune cell PRR to directly bind chitin with high affinity and signal to chitin. Contrary to expectations, oligomers of at least 6 NAGs activated the receptor, whereas oligomers of ≤5 NAGs were inactive or showed antagonistic activity. We thus identified a size-dependent system of immune activation/regulation that appears conserved between plant and animal kingdom. Additionally, interference with chitin-TLR2 binding and signaling provided a way to limit chitin-mediated immune activation and inflammation.

## Results

### Small chitin oligomers activate human and murine immune cells

To illustrate the challenge to study size-dependent aspects of chitin interaction with a cell-surface receptor, we first conducted electron microscopy studies of immune cells incubated with the type of crude chitin used before in the field. These studies showed that crude chitin preparations are not suitable to solve molecular chitin recognition principles since the particles (Expanded View table S1) are as big as entire human macrophages (Fig. 1a) and thus much larger than the size of the NAG chitin subunit and even the known dimensions of typical PRR MAMP-sensing domains (Fig. 1b). This prompted us to re-assess the immunological effects of chitin using pure and endotoxin-free chitin oligomers of defined length (see Methods) comprising 4 to 15 NAGs, which are in the size range of known PRR ligand recognition domains, e.g. TLR ectodomains (Fig. 1b). Whereas fragments with 4 or 5 NAGs (termed e.g. C4 or C5) did not elicit substantial IL-6 and/or TNF release from primary human monocyte-derived macrophages (MoMacs, Fig. 1c) and murine bone marrow-derived macrophages (BMDMs, Fig. 1d) at equimolar concentrations, chitin oligomers of 6 (C6) or 7 (C7) NAGs elicited cytokine release significantly above baseline upon overnight incubation. Preparations of oligomers of 10-15 NAGs (C10-15, MW range 2000-3000 Da, see Supplemental Information) induced cytokine release comparable to the TLR4 agonist LPS. A similar dependence on fragment length ≥6 NAG was observed for the human macrophage-like THP-1 cell line (Fig. 1e). Important control experiments (Expanded View figures S1a,b) showed that the observed effects were not due to endotoxin contamination but rather that chitin oligomers themselves elicited robust immune activation.

**Figure 1:**
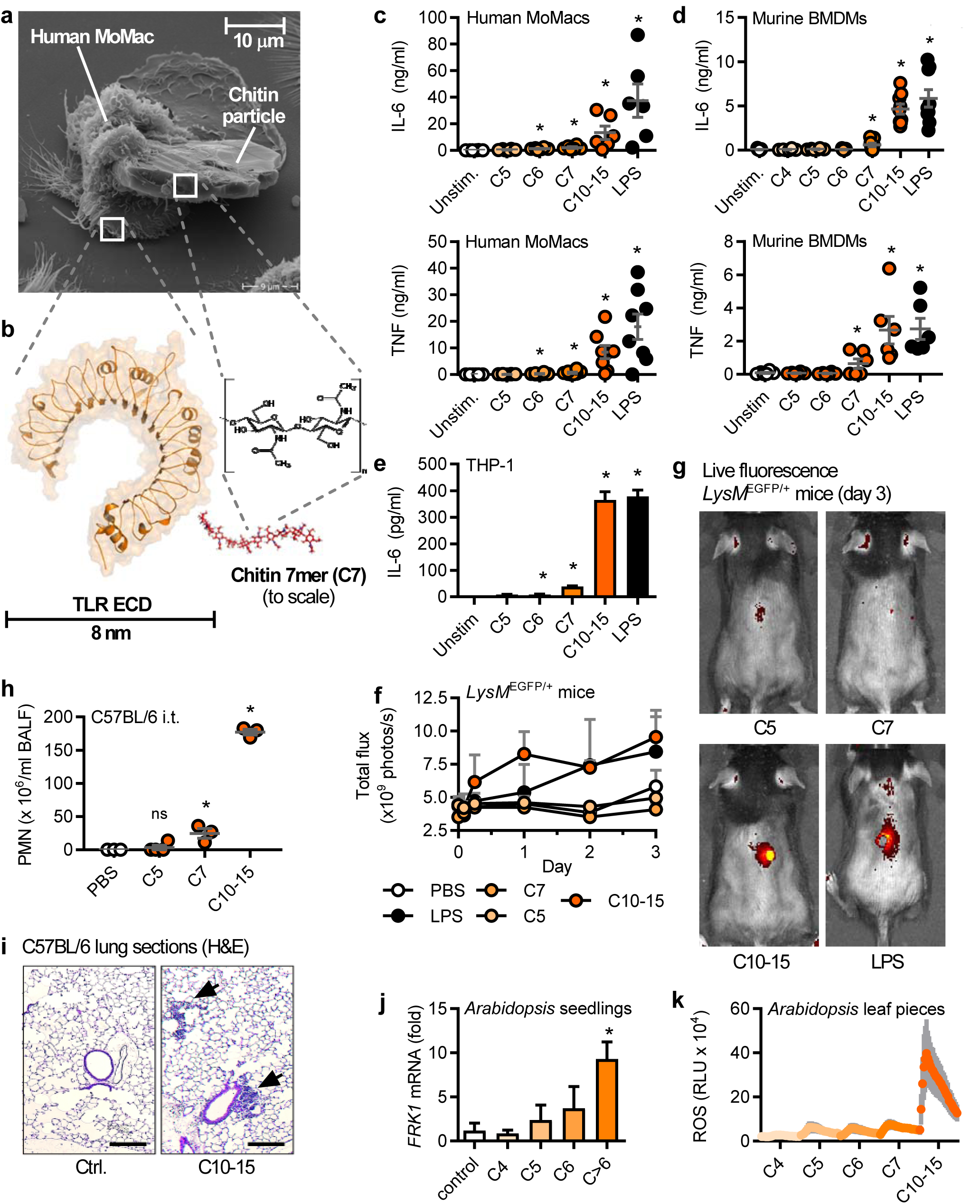
Size-dependent recognition of defined chitin oligomers in humans, mice and plants. (a) Electron micrograph of a human MoMac engulfing a crude chitin particle (n=2). (b) Size comparison (to scale) of typical TLR ectodomain and a chitin 7-mer (seven NAG subunits). Release of IL-6 and/or TNF release (ELISA) from (c) human MoMacs (n=6-8), (d) murine BMDMs (n=6) or (e) THP-1 cells (n=3). (f) GFP *in vivo* fluorescence post intradermal injection of chitin oligomers into *LysM*EGFP/+ C57BL/6 mice (n=3/group). (g) Representative *in vivo* imaging results. Leukocyte infiltration *in vivo* after intratracheal administration assessed by (h) BALF flow cytometry (n=3-4/group) and (i) histology analysis (n=2), scale bar = 200 μm, arrows show cellular infiltrate. *A. thaliana* seedling (j) target gene mRNA induction (n=3) or leaf piece (k) ROS production (0-45 min, n=6). c, d, f, h and k represent data (mean+SEM) combined from ‘n’ (given in brackets for each panel) technical or biological replicates (human donors, mice or plant leaves, respectively). In a, e, g, i and j one representative of ‘n’ (given in brackets for each panel) independent experiments is shown (mean+SD). * p<0.05 according to Wilcoxon signed rank sum (c, d), one-way ANOVA (h) or Student’s t-test (e, j).

To test whether chitin oligomers possessed immunostimulatory properties *in vivo*, several applications were tested in mice. Chitin plays a role in infections and inflammation in both the skin and the lung (Koller et al, 2011; Mack et al, 2015). C10-15 was therefore first injected into the skin of C57BL/6 mice, using a mouse strain expressing enhanced green fluorescent protein (EGFP) under the control of the myeloid-specific promotor, LysM. Thus fluorescently labeled myeloid cells in these mice allow for straightforward *in vivo* imaging of immune activation. C10-15 clearly elicited fulminant local immune cell infiltration indicated by a rapid and strong increase of local EGFP fluorescence (Fig. 1f,g). As chitin has been associated with lung infection and inflammation, C10-15 oligomers were administered intratracheally, leading to a significant influx of white blood cells, especially neutrophils, in bronchio-alveolar lavage fluid (BALF, Fig. 1h) and lung tissue (Fig. 1i). In summary, these results indicated that – contrary to expectations – defined chitin oligomers are immunologically active *in vivo* and *in vitro*, and that immune recognition depends on a minimum number of 6-7 NAG subunits in both human and murine cells. Since size dependent recognition of chitin was reported in plants earlier (Liu et al, 2012), we sought to determine whether this extended across kingdoms. Indeed, application of the same panel of chitin oligomers to *A. thaliana* seedlings was confirmed to elicit innate immune responses such as target gene induction (e.g. *FRK1, MLO12* and *WRKY40*, Figs. 1j and Expanded View figure S1c) and respiratory burst (Fig. 1k) in dependence of oligomer size. Chitin sensing is thus broadly conserved across kingdoms in requiring oligomeric chitin chains of similar chain length for immune activation.

### Oligomeric chitin sensing depends on TLR2

Direct sensing by a typical cell-surface PRR would be expected to lead to a rapid transcriptional response which we indeed observed in THP-1 cells, BMDMs and human MoMacs for *IL6* and *TNF* transcripts (Expanded View figure S2a). Additionally, primary human neutrophils (PMNs) effectively shed CD62L (L-selectin) and released IL-8 within few hours of C10-15 exposure (Expanded View figure S2b). Thus, the time-frame and nature of the elicited response indicated that oligomeric chitin might be sensed by a prototypic PRR expressed by human and murine immune cells. To unequivocally identify this PRR we first tested BMDMs deficient for the TLR adaptor MyD88 and observed a significant reduction in the TNF response to chitin but not the MyD88-independent stimulus, poly(I:C), compared to WT BMDMs (Fig. 2a), indicating the involvement of a MyD88-dependent TLR in chitin sensing. Additionally, TNF release in response to C10-15 as well as the known mycobacterial TLR2 lipopeptide ligand, Pam_3_CSK_4_ (Pam3) was partially reduced in *Tlr2* KO (Fig. 2b) but not *Tlr4* KO BMDMs (*cf.* Expanded View figure S1b). This suggested that at least in mice TLR2 might be involved in molecular chitin sensing. To confirm the involvement of TLR2 in the human system, *MYD88* and *TLR2* were silenced using siRNA in human MoMacs, which led to a reduction in IL-6 release for C10-15 similar to the reduction observed for the TLR2 ligands Pam_2_CSK_4_ (Pam2) and Pam3, but not the TLR4 ligand, LPS, or the TLR8 agonist, R848(Fig. 2c). Blocking of human PMN TLR2 responses, including C10-15, with a well-characterized anti-TLR2 blocking antibody (Meng et al, 2004) but not with a control IgG led to impaired CD62L shedding (Fig. 2d). Furthermore, genetic deletion of *TLR2* in THP-1 cells (Schmid-Burgk et al, 2014) resulted in complete abrogation of IL-6 production in response to C10-15, Pam2 and Pam3, compared to LPS (Fig. 2e). Conversely, genetic complementation of HEK293T cells (which endogenously express TLR5 as well as the known TLR2 co-receptors, TLR1 and TLR6) with TLR2 was sufficient to establish responsiveness to C10-15 in terms of NF-κB activation (Fig. 2f). Of note, Dectin-1, NOD2 and TLR9 reconstitution in HEK293T did not result in chitin responsiveness (Expanded View figure S2c), and *Clec7a* KO (Dectin-1-deficient) immortalized macrophages (iMacs) (Rosas et al, 2008) responded as efficiently to C10-15 as WT macrophages (Expanded View figure S2d). *In vitro*, TLR2 thus was essential for full chitin-mediated cellular activation. To corroborate this in an *in vivo* setting, C10-15 was also applied intradermally or administered intratracheally in WT and *Tlr2 KO* mice. In the skin, WT mice showed significantly higher myeloperoxidase (MPO) activity 12 h post-injection than *Tlr2* KO mice upon treatment with C10-15 but not with LPS, indicating lower PMN activation in *Tlr2 KO* mice (Fig. 2g). Similarly, IL-6 and TNF levels in the BALF of *Tlr2 KO* mice treated with C10-15 were significantly lower than the cytokine levels in WT mice (Fig. 2h), confirming a dependence on Tlr2 *in vivo*. To gain additional evidence for TLR2-dependence of chitin recognition in humans, we analyzed whole blood from healthy individuals without (WT p.753RR) and with heterozygous carriage of a well-known *TLR2* gene variant, rs5743708 (p.753RQ). This variant is strongly associated with pulmonary invasive fungal disease in acute myeloid leukemia patients (OR 4.5, 95% CI 1.4–15.1, *p = 0*.*014*, see (Fischer et al, 2016)) and with up to 40-fold altered cytokine levels during *Candida* sepsis, respectively (Woehrle et al, 2008). Similarly to the mycobacterial TLR2 agonist Pam2, but not to the TLR8 agonist R848, stimulation with C10-15 showed a 100% higher induction of relative *IL10* transcription between 753RR and 753QR allele groups, (Fig. 2i). This was in line with the reported effect of *Candida albicans* TLR2 signaling on IL-10 (Netea et al, 2004) and indicates a shared response of both mycobacterial lipopeptides and fungal chitin via TLR2. Collectively, these results show that TLR2 is a critical mediator of immune activation in response to oligomeric chitin in both humans and mice.

**Figure 2:**
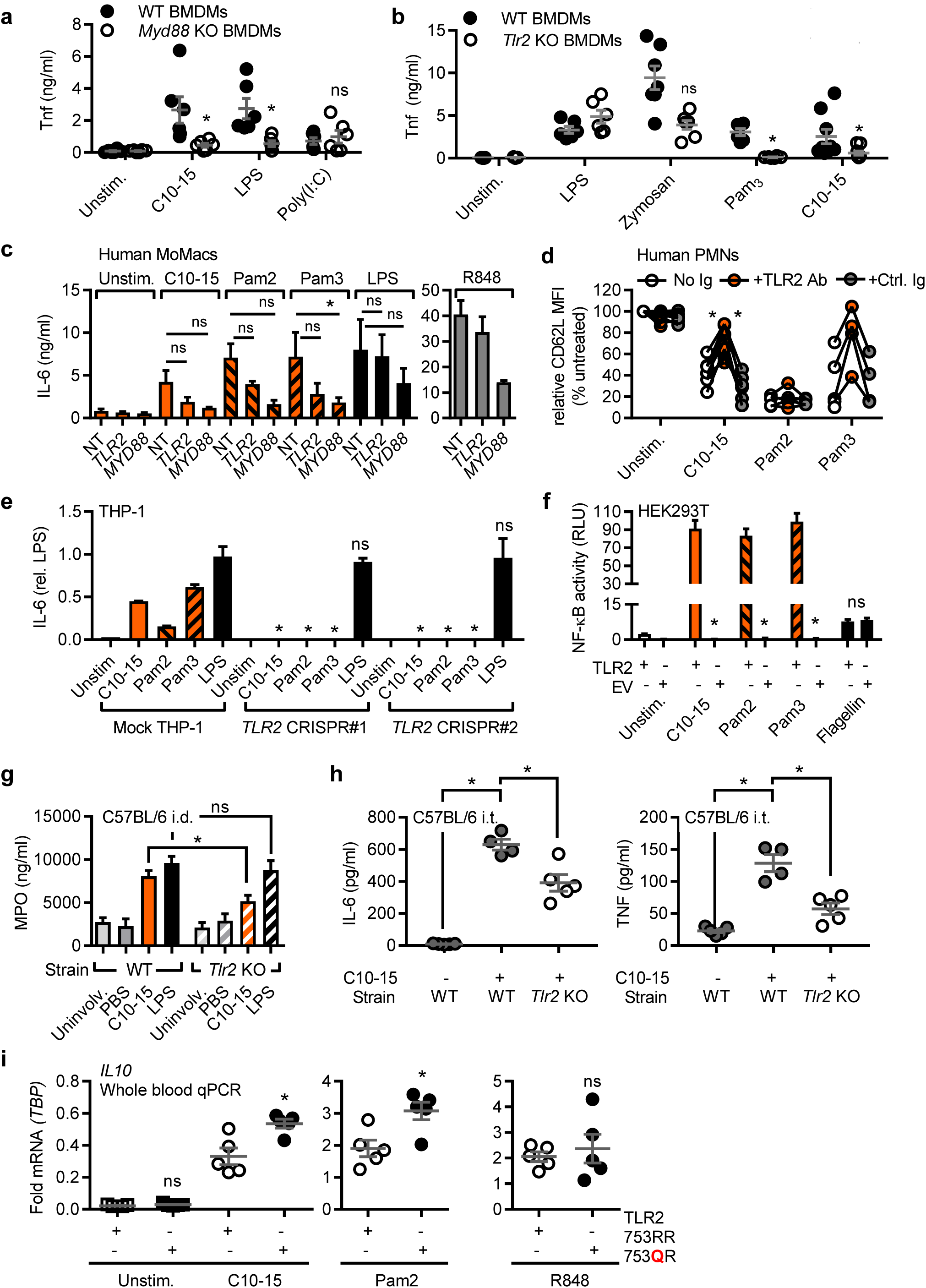
TLR2 is the mediator of chitin oligomer-triggered immune activation. (a-b) TNF released from WT or *Myd88* (a) or *Tlr2* (b) KO mice (total n=6-9/group combined from 3 experiments). (c) IL-6 released from primary MoMacs (n=5) treated with non-targeting (NT), TLR2-or MyD88-specific siRNA. (d) CD62L shedding from human PMNs (n=4-8) with or without isotype control or anti-TLR2 blocking antibody. (e) IL-6 released from mock or *TLR2* Cas9-CRISPR-edited THP-1 cells (n=3). (f) NF-κB activity in empty vector (EV) or TLR2-transfected HEK293T cells (n=3). (g) MPO release from intradermally injected WT or *Tlr2 KO* mice. (h) BALF IL-6 and TNF from intratracheally treated WT or *Tlr2 KO* mice. (i) Relative *IL10* mRNA in whole blood of WT or heterozygous TLR2 R753Q carriers (n=5 each). a, b, c, d, g, h, and i represent data (mean+SEM) combined from ‘n’ technical or biological replicates (human donors or mice, respectively). In e and f one representative of ‘n’ independent experiments is shown (mean+SD). * p<0.05 according to Mann-Whitney U (d, i), Wilcoxon signed rank sum (c), Student’s t-test (a, b, e, f, g) or one-way ANOVA with Dunnett’s multiple comparison (h).

### Oligomeric chitin is a unique fungal TLR2 agonist

Several earlier *in vitro* and *in vivo* results had implicated TLR2 in fungal infection (reviewed in (Cunha et al, 2010)) but the precise molecular nature of the fungal TLR2 agonist has so far remained unclear. This is epitomized by the finding that the complex *Saccharomyces cerevisiae* cell wall preparation, zymosan, is well-known to possess TLR2-stimulatory activity (Gantner et al, 2003) but despite >1,200 published immunological studies using zymosan, the molecular nature of this ‘TLR2 activity’ is unknown. Interestingly, TLR2 (but not Dectin-1) activity can be “depleted” by hot alkali treatment. Thus, among the known components of fungal cells that are chemically sensitive to hot sodium hydroxide treatment, we noted that chitin was previously chemically de-acetylated by such treatment (No et al, 2000). Interestingly, hot alkali depletion of both zymosan and C10-15 led to reduced NF-κB activation in TLR2-transfected HEK293T cells (Fig. 3a). ‘Depleted’ zymosan also showed reduced TLR2-dependent IL-8 release from primary PMNs but rather increased ROS production (Fig. 3b). ROS production was not detectable by known TLR2 agonists like Pam2 or Pam3 and is thus TLR2-independent. Furthermore, mass spectrometry confirmed the de-acetylation of chitin upon ‘depletion’ and that soluble material released by *in vitro* treatment of insoluble zymosan with a recombinant bacterial chitinase contained di-NAG (Expanded View figure S3a) in line with earlier reports (Di Carlo & Fiore, 1958). Our results point to chitin as a relevant TLR2-stimulatory component of zymosan, identifying the molecular moiety that links zymosan, fungal infection in general and the role of TLR2.

**Figure 3:**
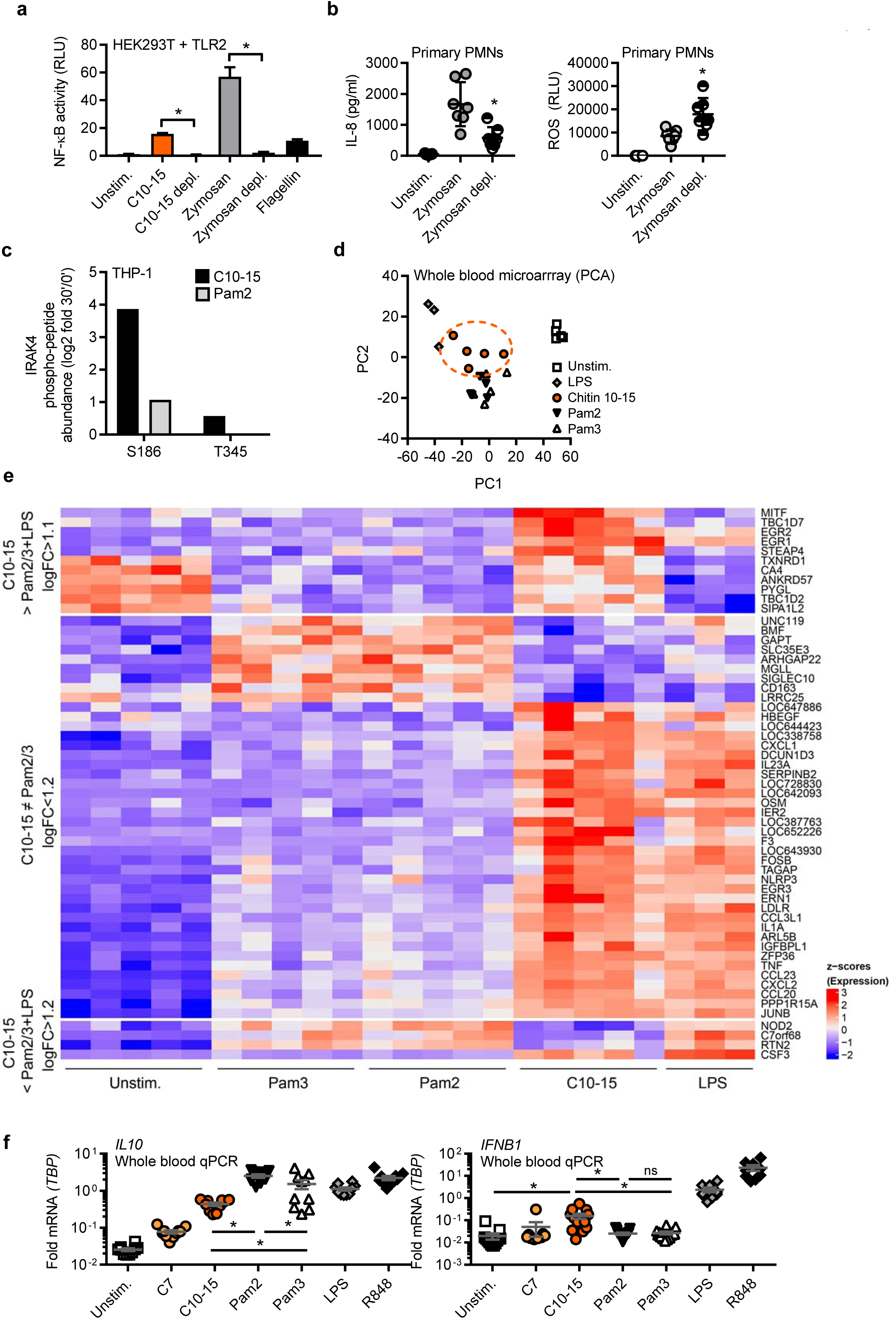
Chitin is a unique fungal TLR2 ligand. (a) NF-κB activity in TLR2-transfected HEK293T cells. (n=3). (b) IL-8 release and ROS production from primary PMNs (n=7). (c) IRAK4 S186 and T345 phospho-site containing IRAK4 peptides quantified in proteome-wide phospho-screen in THP-1 cells (n=1). Whole blood principal component analysis (d) and intensities (e) of microarrays (selected for log fold changes, logFC, as indicated and grouped by induction of C10-15 relative to other treatments) from healthy donors (n=3-5). (f) Relative cytokine mRNAs in whole blood stimulation (n=5-13). b, d, e and f represent data (mean+SEM) from ‘n’ technical or biological replicates (donors). In a and c one representative of ‘n’ independent experiments is shown (mean+SD). * p<0.05 according to Wilcoxon signed rank sum (b, f) and Student’s t-test (a); for c, d see Expanded View section.

We next wondered whether chitin-elicited, i.e. fungal, TLR2 signaling further differed from mycobacterial lipopeptide TLR2 signaling at the levels of signal transduction and transcriptional profiles under equimolar ligand concentrations. First, mapping of a dataset from a preliminary phospho-proteomics screen (dimethyl-labeling global phospho-proteomics, see Methods) in THP-1 cells to KEGG pathways indicated that within 30 minutes C10-15 activated signaling pathways typically engaged by cell surface TLR ligands in macrophages, e.g. MAPK, PI3K and NF-κB ((Weintz et al, 2010) and Expanded View table S2). We also compared specific phospho-peptides in Pam2 and C10-15 stimulated (30 min at equimolar concentrations) samples. Although the majority of phosphorylation events were shared between Pam2 and chitin, certain phospho-peptides mapping to MAPK pathway members, e.g. MEK2, cPLA2, STMN1, MAPKAPK, and PAK1/2, were more abundant upon C10-15 stimulation than after treatment with Pam2 (Expanded View table S2). Additionally, a novel phosphorylation site at Ser 186 in the TLR2 pathway kinase IRAK4 was identified and found in 16-fold greater abundance in C10-15-stimulated vs. 2-fold in Pam2-stimulated samples, compared to unstimulated samples (Fig. 3c), hinting that C10-15 downstream signaling may have certain distinct features compared to other TLR2 ligands. To investigate this more broadly, we used microarrays to study how chitin-stimulation impacted the transcriptomic profiles in whole blood from healthy donors, compared to Pam2 and Pam3 stimulation. Principal component analysis (PCA, Fig. 3d, see Supplemental Methods and Expanded View table S3 and Expanded View code) showed that in the projection of PC1, which contained cytokine genes typically regulated by TLRs (e.g. *IL6 or CCL3*), C10-15 and Pam2 and Pam3 were highly similar, suggesting a considerable overlap of regulated genes as could be expected by ligands sharing the same receptor (Expanded View figure S3b). However, in PC2 (loaded by more IFN-related genes such as *IFNB1, IFIT1, IFIT3, OASL, CXCL11*), C10-15 was also distinct from Pam2/Pam3 (Fig. 3d, Expanded View figure S3b), illustrating a distinct activation pattern of gene regulation for C10-15 compared to the Pam2/3 TLR2 ligands in blood cells. Excerpts of a clustering analysis showed distinct groups of genes uniquely up- or down-regulated for C10-15 compared to Pam2/Pam3 and/or LPS (Fig. 3e): Selected genes upregulated significantly >2-fold (log2 fold change>1.1) more by C10-15 compared to Pam2, Pam3 and LPS included the transcriptional regulators, early growth response (*EGR*) 1 and 2 (top block of Fig. 3e); genes upregulated >2-fold (log2FC>1.2) more by both C10-15 and LPS compared to Pam2/3 included genes such as *EGR3*, several chemokines and the cytokines *IL1A, TNF* and *IL23A*, as well as the PRR *NLRP3* (middle block of Fig. 3e); on the other hand, C10-15 downregulated (log2FC>1.2) genes like the PRR *NOD2* more than any other treatment (lower block of Fig. 3e). Validation by qPCR in additional donors (Fig. 3g and S3b) confirmed that chitin induced typical TLR-regulated cytokine genes. However, depending on the gene of interest, at equimolar ligand concentrations the strength of induction differed between the TLR2 ligands, similar to the microarray. For example, C10-15 induced less *IL10* (3.5 or 5.7-fold mean transcript levels) but significantly higher levels of *IFNB1* (5.8 or 6.2-fold) compared to Pam2 or Pam3 (Fig. 3f). These results indicate that at equimolar concentrations, oligomeric chitin acts as a novel TLR2 agonist with largely overlapping but also certain distinct (e.g. *IFNB1* > *IL10* induction) properties compared to known mycobacterial TLR2 agonists. Potentially, these differences may be attributable to (a) co-receptor(s) that functions differently than TLR1 or TLR6 for mycobacterial ligands.

### The TLR2 ectodomain directly binds to chitin in solution and on the fungal cell wall

Given that all other previously known TLR2 agonists are lipopeptides (Jin et al, 2007; Kang et al, 2009), the finding that TLR2 senses the oligosaccharide chitin was initially surprising and warranted further confirmation by direct binding assays. In a flow cytometric setup, Alexa647-labeled C10-15 interacted with recombinant murine TLR2 ectodomain-human IgG1-Fc fusion protein (mTLR2-Fc) but not with a control IgG1-Fc (Figs. 4a). Additionally, microscale thermophoresis (MST, see (Jerabek-Willemsen et al, 2011)) demonstrated a dose-dependent binding of Alexa647-C10-15 to murine (Figs. 4b) and human (Expanded View figure S4a) TLR2 ectodomains with Kd values in the low nanomolar range, namely 6.65 +/-1.69 nM and 7.18 +/-0.3 nM, respectively. This binding was dependent on correct folding of the TLR2 ectodomain since controlled heat-denaturation of the TLR2 protein (Jimenez-Dalmaroni et al, 2015) abrogated C10-15 binding completely. Thus, isolated chitin oligomers directly bound to the TLR2 ectodomain *in vitro*. TLR2 also bound to *C. albicans* cells: when stained with mTLR2-Fc or control IgG1-Fc in combination with anti-Fc-Alexa594, *C albicans* cells assessed by flow cytometry showed a higher median fluorescence intensity for mTLR2-Fc-than for control IgG-stained samples (Fig. 4c), indicative of TLR2 binding. The control stain, calcofluor white (CFW), which broadly stains chitin-rich cell wall areas, showed comparable staining across samples. In confirmation of this result, fluorescence microscopy showed few but distinct mTLR2-Fc-stained areas within the cell wall of stained *C. albicans* (Fig. 4d). These data suggest that the TLR2 ectodomain can bind directly to chitin in the fungal cell wall but potentially with few accessible binding sites.

**Figure 4:**
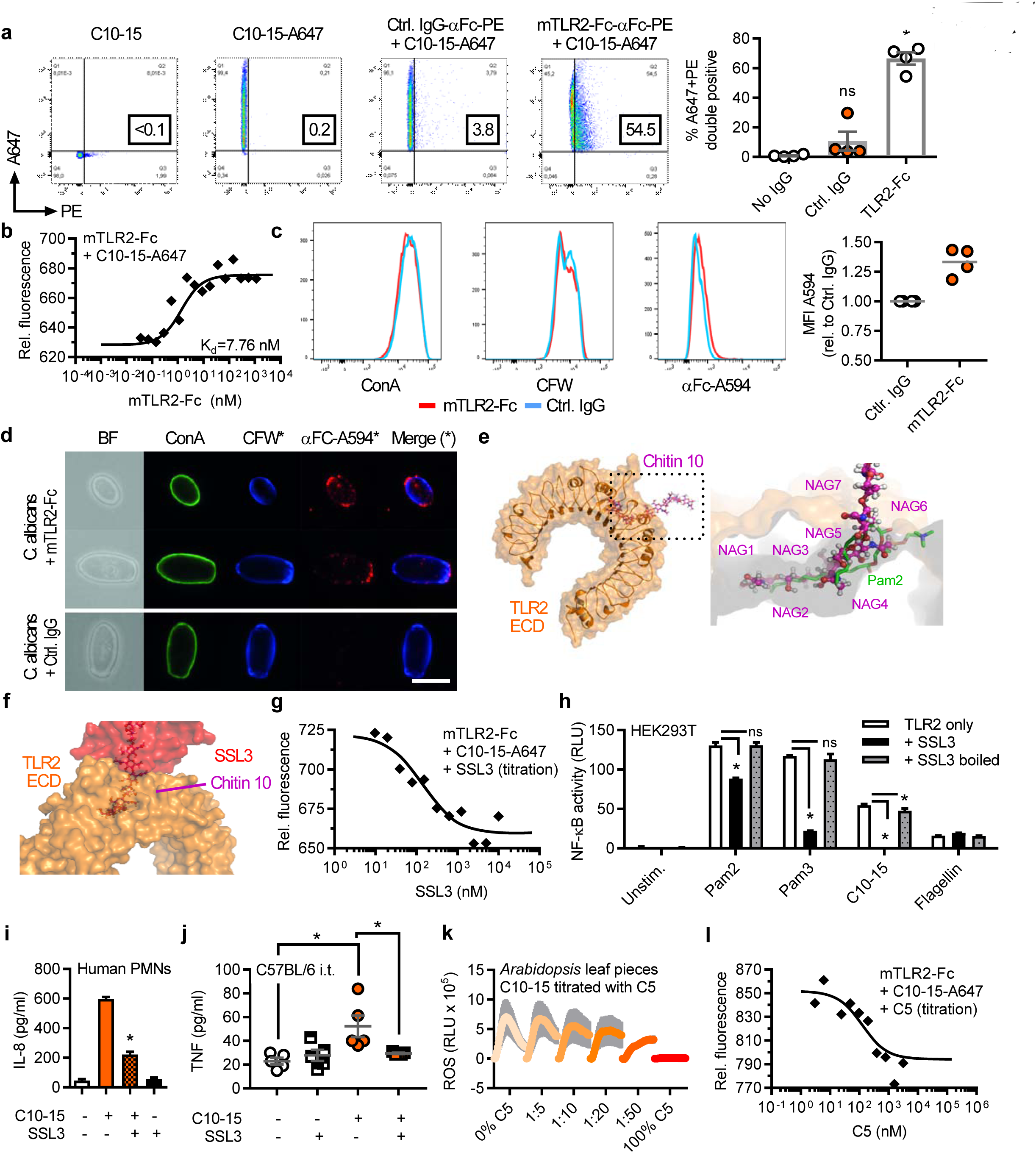
Direct chitin TLR2 binding and immune activation can be blocked by SSL3 or short chitin oligomers. Flow cytometric quantification (a) or microscale thermophoresis (MST) analysis (b) of Alexa647-labeled C10-15 interaction with mTLR2-Fc Protein (n=4 each). (c) Flow cytometry and (d) fluorescence microscopy of *Candida albicans* stained with control IgG or mTLR2-Fc anti-Fc-Alexa594, together with ConA-Alexa488 and CFW (n=4 each). Scale bar 5 μm. (e) Docking of chitin 10 (magenta) into the TLR2 (orange) Pam2 (green) lipopeptide binding pocket (close-up, pdb: 2z7x), (f) overlayed with SSL3 from a SSL3-inhibited TLR2 complex (pdb: 5d3i). (g) MST analysis of Alexa647-labeled C10-15 and mTLR2-Fc protein with SSL3 titration (n=2). (h) NF-κB activation in TLR2-transfected HEK293T cells (n=3) or (i) IL-8 release from primary PMNs (n=2) without or with SSL3. (j) BALF TNF in C57BL/6 mice (n=5/group) upon C10-15 administration without or with SSL3. (k) ROS production between 0 and 45 min post chitin application in *A. thaliana* leaf pieces (n=6). (l) MST analysis of Alexa647-labeled C10-15 and mTLR2-Fc Protein with C5 titration (n=2). a, c, k and j represent data (mean+SEM) combined from ‘n’ biological replicates (human donors, mice or plant leaves, respectively). In a, b, c, d, g, h, i and l one representative of ‘n’ independent experiments is shown (mean+SD). * p<0.05 according to Mann-Whitney U (a, b, d, e), Student’s t-test (h, f, m, i) or one-way ANOVA with Dunnett’s multiple comparison (j).

Crystallographic evidence shows that mycobacterial TLR2 ligands are bound in a hydrophobic TLR2 pocket (Jin et al, 2007; Kang et al, 2009). *In silico* docking studies (see Methods) indicated that 5 NAG subunits of chitin, a relatively apolar molecule (Bueter et al, 2011; Bueter et al, 2013), can be accommodated in the TLR2 hydrophobic pocket (Fig. 4e and Movie S1). Mycobacterial lipopeptide binding and TLR2 activation can be blocked by the *S. aureus* immune evasion protein staphylococcal superantigen-like protein 3 (*SSL3*) (Koymans et al, 2015). Thus, if chitin is bound via the same binding pocket as lipopeptides, SSL3 would be expected to impair chitin TLR2 binding and consequently immune activation (Fig. 4f). Indeed, SSL3 decreased C10-15-TLR2 binding in a dose-dependent manner (Fig. 4g) and reduced NF-κB activation in TLR2 HEK293T cells and IL-8 release from primary neutrophils(Fig. 4h,i). Binding of chitin to the hydrophobic pocket was further supported by the comparable effects of the known TLR2 blocking antibody on both chitin and lipopeptides (*cf.* Fig. 2d), the effect of TLR2 point mutants narrowing the hydrophobic pocket (see Expanded View figure S4b-d) and additional molecular modeling (Expanded View figure S4e-h, Movie S2). Thus chitin-TLR2 binding appears to involve the free ends of at least 5 NAG long oligomers, which explains why fungal cells, where chitin is polymeric and cross-linked to other cell wall components, only have limited numbers of accessible TLR2 epitopes (*cf*. Fig. 4c,d). Since chains of only 5 NAGs were not sufficient for signaling (*cf.* Fig. 1c-e), we speculate that NAG subunits protruding from the binding pocket (NAG=6) are essential for making additional contacts with another TLR2 ECD or a co-receptor, which may lead to receptor dimerization events known to establish TLR2 signaling for other ligands (Jin et al, 2007; Kang et al, 2009). For fungal infections, the effect of *S. aureus* SSL3 on fungal chitin sensing is noteworthy since co-infections of *S. aureus* and pathogenic fungi such as *C. albicans* frequently occur (Morales & Hogan, 2010).

### Interference with chitin-TLR2 binding suppresses chitin-mediated inflammation

Since we identified TLR2 as the direct mediator of inflammatory signaling in response to chitin, chitin-TLR2 binding would represent a critical regulatory point in both the fungal immune response and for therapeutic intervention in chitin-mediated pathologies such as lung inflammation. Having observed that SSL3 was a potent antagonist of chitin binding and receptor activation *in vitro*, we wondered whether blocking chitin-TLR2 binding might also reduce immune activation and inflammation in the *in vivo* model of lung inflammation. Application of SSL3 together with C10-15 indeed significantly reduced both TNF and IL-6 levels in the BALF of intratracheally treated animals compared to C10-15 alone (Figs. 4j and S5a). An alternative mode of blocking chitin binding, was gleaned from earlier studies in plants (Liu et al, 2012) and confirmed here, namely that short (<6) NAG chains reduce immune activation elicited by longer NAG chains (Fig. 4k). Based on these and our modeling data (*cf.* Fig. 4, S4), we speculated that inactive chitin fragments such as C5 are probably too short to bridge receptors to initiate signaling but may nevertheless compete with stimulatory C10-15 for the TLR2 binding pocket. Indeed, C5 dose-dependently inhibited the binding of a constant amount of Alexa647-chitin to TLR2 in MST measurements (Fig. 4l), suggestive of a potential inhibitory effect of chitin oligomers of ≤5 NAG chain length. In preliminary experiments in THP-1 cells and primary MoMacs an inhibitory effect on IL-6 release was indeed observed when the cells were pre-treated with C5 before the addition of stimulatory C10-15 (Expanded View figure S5b, c). Collectively, our data indicate that chitin binding to TLR2 is direct and, depending on oligomer size may promote or even restrict immune activation. Additionally, receptor binding emerges as a physiologically relevant and therapeutically attractive step in chitin-mediated immune activation and amenable to interference by different TLR2-antagonistic strategies.

## Discussion

Although different receptors have been hypothesized to mediate the pleiotropic pathological effects of chitin in asthma and fungal infection, so far there have been few molecular studies and the identity of a binding receptor on immune cells has remained elusive, owing largely to the unavailability of defined chitin ligands. We here identify oligomeric chitin as a high affinity, direct and novel TLR2 ligand, activating multiple innate cell types (Macrophages, DCs and PMNs) in a size-dependent manner. Several important points warrant further discussion:

Firstly, by using chitin oligomers of defined lengths our study now makes a firm mechanistic connection between a (i) vast number of non-mechanistic chitin studies, (ii) chitin-related pathologies and (iii) TLR2 a key responsible PRR. Although a lung chitin phenotype for *Tlr2* KO was reported before (Da Silva et al, 2008), other PRRs (e.g. TLR9 or NOD2) were equally proposed as chitin receptors (e.g. (Wagener et al, 2014)). Unfortunately, all previous studies had a non-mechanistic approach, used crudely purified chitin and did not show direct binding, which left open whether the observed effects were direct. Our study now dispels these ambiguities and provides unequivocal experimental evidence for TLR2 as a molecular chitin immune receptor. Our results now provide a missing molecular link between fungal infection and the previously observed dependence on TLR2 in murine fungal infection models (Cunha et al, 2010; Goodridge & Underhill, 2008) and, importantly, in patients with the functional rs5743708 (p.753RQ) *TLR2* allele (Fischer et al, 2016; Woehrle et al, 2008). Of note, the effect of SSL3 on chitin sensing warrants further analysis in *S. aureus/C. albicans* co-infections (Morales & Hogan, 2010): Based on our *in vitro* and *in vivo* results, the SSL3 counter-strategy employed by *S. aureus* to tune down the effect of its TLR2-activating lipoteichoic acid MAMPs (Koymans et al, 2015), may not only lead to a dampened host response against *S. aureus* but concomitantly also impair the chitin-TLR2-mediated anti-fungal response.

Mechanistically, our data indicate that TLR2 is the first TLR to bind multiple classes of MAMP biomolecules, namely both bacterial lipopeptides and fungal oligosaccharides. Although our data firmly indicate TLR2 to bind and be involved in signaling to chitin, they leave open the possibility of (a) co-receptor(s) contributing to full activation. The differences observed between lipopeptide and chitin signaling and gene regulation may indicate this does not involve the canonical TLR2 co-receptors TLR1 or TLR6 and warrant further research in the future. The concept that TLR2 has a preference for free ends of ≥6 NAG, both as soluble oligomers or in the context of the fungal cell wall, warrants refinements (Expanded View figure S6) of previously proposed concepts of size dependent chitin-based fungal recognition which focused on the macromolecular (insoluble) range (Da Silva et al, 2009; Lee et al, 2008). Although insoluble large chitin fragments are also detected by TLR2 when fungal pathogens are directly encountered by myeloid cells (*cf.* Fig. 4c,d) human chitinases (e.g. the endochitinase CHIT1 endochitinase, (Mack et al, 2015; Stockinger et al, 2015)) would not only create additional free chitin ends required for TLR2 sensing but also soluble, smaller fragments that could be detected by TLR2 on distal immune cells for a full anti-fungal immune response. In a manner analogous to the action of LPS binding protein and CD14 as ‘shuttles’ for LPS, chitinase-like proteins may augment chitin fragment solubility, stability and presentation to TLR2. As fungal pathogens are cleared, further chitinase activity may eventually generate antagonistic fragments with ≤5 NAGs, that may compete with stimulatory fragments and thus be part of a negative feedback loop to restrict excessive inflammation. Such degradation-based immune control has already been described, e.g. for peptidoglycans in *Drosophila* (Wang et al, 2006) or hyaluronan in humans (Scheibner et al, 2006). However, successive degradation has been seen to only lead to the eventual loss of agonistic MAMPs. Our study points to the possibility of a downstream anti-inflammatory loop based on small antagonistic MAMP fragments to also operate in humans and mice, which so far has only been reported for plants (Liu et al, 2012). It could be speculated that recognition based on a minimum size of chitin (≥6 NAG) might potentially also act as a safe-guard against immune activation by NAG mono-or disaccharides that might arise during host N-linked core oligosaccharide synthesis.

Finally, from a translational perspective, our study raises the possibility that chitin oligomers could serve as tools for developing therapies in chitin-mediated inflammatory disease conditions, such as fungal- or house-dust mite-related asthma and lung fibrosis (Van Dyken et al, 2017). Efforts in the latter area could now center on chitin-TLR2 binding and activation as critical patho-biological events, and on developing biologicals (e.g. SSL3-derivatives or blocking antibodies), short chitin oligomers or other ‘chito-mimetic’ TLR2 antagonists structurally resembling short-chain NAGs to block these steps. Such targeted approaches may bring therapeutic benefit to the vast and growing number of patients suffering from allergic asthma (To et al, 2012). Therapeutic opportunities in the treatment or control of some of the millions of cases of fungal infections worldwide (Brown et al, 2012) could also be envisaged.

## Methods

### Reagents and antibodies

All chemicals were from Sigma unless otherwise stated. Polymeric crude chitin and C4, C5, C6, C7 chitin oligomers and C10-15 chitosan oligomers were from Isosep, Elicityl and Carbosynth, respectively. Chitin 10-15 was generated from chitosan oligomers by acetylation (Bueter et al, 2011) Purities of >95%, acetylation of >90% and coupling to Alexa647 were achieved. Further details and all other reagents are described in Supplemental Information.

### Plasmid constructs

Plasmids for Flag-tagged TLR2 and TLR9 were from I. Bekeredjian-Ding and A. Dalpke (Medical Microbiology, Heidelberg University, Germany). Plasmids expressing human NOD-2 were from T. Kufer (Hohenheim University, Germany) or for human Dectin-1 from G. Brown (Aberdeen University, UK). TLR2 hydrophobic pocket mutants were generated using the QuikChange II XL site-directed mutagenesis kit (Agilent) according to the manufacturer’s instructions using primers shown in Expanded View table S4 and verified by automated DNA sequencing (GATC Biotech).

### Growth and fixation of *Candida albicans*

*C. albicans* SC5314 was grown in synthetic complete media (Formedium SC broth 2% glucose, additionally supplemented with 25 mg/L adenine sulfate) by shaking at 150 rpm at 30°C overnight, fixed by adding paraformaldehyde to 3.7% for 1 hour and harvested by centrifugation and washing in sterile Dulbecco’s phosphate-buffered saline (DPBS).

### Analysis of TLR2 binding to *Candida albicans* cells

1x106 fixed *C. albicans* cells were incubated with 1 μM recombinant mouse TLR2 Fc chimera protein or corresponding control human IgG1 Fc overnight. Next day cells were stained with Alexa594-conjugated mouse anti-human IgG (Jackson Immunoresearch 209-585-098), 25 μM Calcofluor White and 50 μg/mL Concanavalin A CF488A (Biotum 20967 and 29016)at room temperature in DPBS. For microscopy, cells were imaged on Poly-L-Lysine coated coverslips on a Nikon Eclipse Ti2 system at 100x magnification. Flow cytometry was conducted with a BD LSRFortessa^™^ system. Further details in Expanded View section.

### Flow cytometric analysis of TLR2 chitin interaction

0.5 μM Alexa647-labeled C10-15 was incubated with 70 nM of recombinant mouse TLR2 Fc chimera protein (R&D 1530-TR-050) or a corresponding control IgG1-Fc in 200 μl DPBS over night at 4°C, stained with 1:15 diluted human Fcγ-specific PE-conjugated F(ab’)2 fragments (Jackson Immunoresearch 109-116-098) and measured on a BD FACSCanto^™^ || system with 488 and 633 nm laser excitation for the respective fluorophores. Further details see Supplemental Information.

### Microscale thermophoresis analysis of TLR2 chitin interaction

Microscale thermophoresis (Jerabek-Willemsen et al, 2011) was conducted with fluorescence-labeled chitin and recombinant TLR2. In brief, 1:1000 diluted Alexa647-labelled C10-15 chitin (c=17.7 μM) was mixed 1:1 with 2-fold serial dilutions of mTLR2-Fc or hTLR2 (R&D Systems, c = 1150 nM or 1900 nM maximal concentrations in PBS, see Supplemental Information), incubated in the dark at RT overnight with gentle shaking. Solutions were then transferred to Nanotemper capillaries and measured on a Monolith NT.115 instrument and dataset with the highest ΔF fitted using Nanotemper software.

### Dual NF-kB luciferase assay and immunoblot in HEK293T cells

HEK293T were transfected with 25 ng TLR2 plasmid, 10 ng TLR9 plasmid, 0.25 ng NOD2 plasmid or 50 ng Dectin-1 plasmid or empty vector (EV) backbone, firefly luciferase under the NF-κB promoter (100 ng) and constitutive Renilla luciferase reporters (10 ng), stimulated for 18 h with the respective PRR agonists (MDP 200 nM, zymosan 100 μg/ml, CpG, Pam2 or 3 and C10-15 at 1 μM) and DLA measured. For immunoblot equal protein lysates in RIPA buffer were run on 8% Tris-Glycine gels, separated, blotted on nitrocellulose membrane and probed with anti-Flag (Sigma F7425) and anti-rabbit IgG-HRP (Vector laboratories PI-1000) using a BlotCycler (Biozym) and a Vilbert Lourmat CCD system for ECL detection. Further details see Supplemental Information.

### ELISA

Cytokines were determined in half-area plates (Greiner Bio-One) using triplicate points on a standard plate reader. The assay was performed according to the manufacturer’s suggestions (Biolegend), using appropriate (1/3 – 1/20) dilutions of the supernatants.

### Respiratory burst (ROS) assay and cytokine analysis from Dectin-1-deficient macrophages

Immortalized murine macrophages from *CLEC7A* KO (Dectin-1 deficient) and the respective WT mice were originally described (Rosas et al, 2008) and were a kind gift of P. Taylor (Cardiff University, UK). ROS or cytokine release upon PRR agonist addition (C10-15 and Pam2 at 5 μM and zymosan at 50 μg/ml) were measured as described in Expanded View section.

### THP-1 cell line analysis

The human monocytic THP-1 cell line (Invivogen) and two THP-1-based TLR2 Cas9-CRISPR-edited clones or their parental cell lines (a gift from V. Hornung, Gene Center, Munich, Germany (Schmid-Burgk et al, 2014) differentiated with 300 ng/ml PMA overnight were rested for 48 hours without PMA and stimulated with C7, C10-15, Pam2, Pam3 at 5 μM, LPS at 1 μg/ml, or zymosan at 25 μg/ml. For competition experiments, C10-15 was used at 5 μM and C5 at 0.2, 1 and 5 μM and pre-incubated for 2 h. qPCR of mRNA induction relative to TBP using gene-specific TaqMan primers were performed as described in Supplemental Information (ThermoFisher, Expanded View table S5) or ELISAs as described before.

### Dimethyl-labeling of THP-1 cells and global phosho-proteome mass spectrometry analysis

THP-1 cells (Invivogen) were prepared and stimulated as before with PBS, 5 μM C10-15 or 5 μM Pam2 for 30 minutes. Dimethyl-labeling of equal amounts of lysate, sample workup, MS acquisition, peak annotation and database searches was done as described in Expanded View section.

### Human study subjects and sample acquisition

All healthy blood donors included in the analyses of immune cells for this study provided their written informed consent before study inclusion. Approval for use of their biomaterials was obtained by the local ethics committee at the University of Tübingen, in accordance with the principles laid down in the Declaration of Helsinki. Buffy coats were obtained from blood donations of healthy donors were received from the Center for Clinical Transfusion Medicine (ZKT) at the University Hospital Tübingen and whole blood from voluntary healthy donors was obtained at the University of Tübingen, Department of Immunology.

### Mice

*Myd88* and *Tlr2 KO* mice (both on a C57BL/6 background, originally a gift from H. Wagner, Ludwigs-Maximilian University, Munich) and C3H/HEJ (*Tlr4*^*LpsD*^, Jackson) were used between 8 and 20 weeks of age in accordance with local institutional guidelines on animal experiments and under specific locally approved protocols for sacrificing and *in vivo* work. All mouse colonies were maintained in line with local regulatory guidelines and hygiene monitoring.

### Isolation and analysis of primary human monocyte-derived macrophages

Primary monocytes were isolated from heparinized whole blood (Department of Immunology) using standard Ficoll density gradient purification andanti-CD14 magnetic bead positive selection (Miltenyi Biotec, >90% purity assessed by anti-CD14-PE flow cytometry). For macrophage differentiation, cells were grown with GM-CSF (Prepro Tech) for 6 days, re-seeded and treated with LPS (100 ng/ml), C10-15, Pam2 and Pam3 (all 5 μM), R848 (5 μg/ml). For competition experiments, C10-15 was used at 5 μM and C5 at 0.2 μM pre-incubated for 2 h. For electron microscopy (performed by J. Berger, Max-Planck-Institute, Tübingen, Germany) cells were seeded on poly-L-lysine-treated cover slips in 24 well plates. Specimen preparation and EM analysis was done according to standard procedures. For RNAi experiments, MoMacs were transfected with 35 nM of corresponding siRNA (see Expanded View table S6, GE Dharmacon) using Viromer Blue (Biozol) on day 5 of GM-CSF differentiation. On day 6 cells were stimulated and supernatants harvested on day 7. Further details see Supplemental Information.

### Isolation and analysis of primary human neutrophils

Primary neutrophils were isolated by Ficoll density gradient purification using ammonium chloride erythrocyte lysis to >97% purity and no signs of pre-activation. Where appropriate, the cells were pre-treated with a purified anti-mouse/human anti-CD282 (TLR2) Ab (Biolegend 121802, (Meng et al, 2003)), a Sigma anti-HA (H9658) isotype control or SSL3 for 30 min and subsequently stimulated with the PRR agonists C10-15 (5 μM), PMA (500 ng/ml), LPS (200 ng/ml), Pam2 and Pam3 (0.5 or 5 μM), zymosan and zymosan depleted (both 25 μg/ml) for 1 h (CD62L shedding using anti-CD62L-BV421) or 4 h (IL-8 ELISA). For ROS analysis, cells were incubated with stimuli together with DCF at 10 μM and fluorescence measured continuously for 3 h. Further details see Supplemental Information.

### Human whole blood microarray and qPCR analyses

For whole blood assays 1 ml freshly drawn heparinized blood was aliquoted and stimulated for 3 h. Subsequently, plasma was removed and erythrocyte lysis performed. The remaining cells were frozen at -80 °C in RLT buffer for RNA isolation (QIAamp RNA Blood Mini Kit, Qiagen), genomic DNA digestion (Turbo DNA-free Kit, ThermoFisher), reverse transcription to cDNA (High Capacity RNA-to-cDNA Kit; ThermoFisher) and analysis by qPCR using gene-specific TaqMan Primers (Thermo, see Expanded View table S5) or microarray (Illumina HumanHT-12 v4.0 Expression BeadChip Kit on an Illumina HiScan array scanner). Microarray expression data was analyzed with R, as summarized in the Expanded View Methods (including R code producing the figures and further insights into the data as described therein).

### Isolation and analysis of mouse bone marrow derived macrophages (BMDM)

Mice were sacrificed, femurs and tibia were then opened and bone marrow (BM) cells flushed out passed through a 0.22 μM strainer and differentiated with GM-CSF for 6 days. The resulting BMDMs were seeded in 96 well plates and TNF and/or IL-6 levels were measured after 18 h incubation with the following stimuli: C5 (5 μM), C6 (5 μM), C7 (5 μM), C10-15 (5 μM), LPS (1 μg/ml), poly(I:C) (20 μg/ml), Pam2 (5 μM), Pam3 (5 μM), zymosan and zymosan “depleted” (both 10 μg/ml). For measuring ROS activity 75.000 cells were seeded and assayed as described above for WT and *Clec7a*-deficient immortalized macrophages.

### *In vivo* dermal inflammation analysis

For female LysM-EGFP mice expressing EGFP in myeloid cells (Cho et al, 2012), the dorsal backs were shaved under 2% isoflurane, injected intradermally (i.d.) with chitin oligomers C10-15, C7 and C5 at 5 μM concentration in PBS, 1 mg/kg LPS or PBS alone, and imaged *in vivo* on a Lumina III IVIS (PerkinElmer) as described in Supplemental Information. 10 mm punch biopsies were also taken from the injection site at 3 days after treatment, weighed, then homogenized in protein lysis buffer for Myeloperoxidase assays (MPO; R&D Systems).

### *In vivo* lung inflammation analysis

WT and *Tlr2* KO mice were anaesthetized using ketamine-xylazine solution and a vertical cut was made on the neck to expose trachea to instill 30 μl of 1 mM chitin oligomers using a Hamilton syringe. For competition experiments with SSL3, 25 μl of an endotoxin-free 50 μg/ml (2.6 mM) of SSL3 solution was first instilled into the trachea followed by instillation of 30 μl of a 1 mM C10-15 solution. The wound was afterwards sealed using Vetbond tissue adhesive. Mice were euthanized 12 h post instillation of chitin to harvest BALF. Lung tissues were inflated at 1 atm pressure with 0.5% low melting point agar and fixed using formalin. Lung sections were embedded and sections were stained with hematoxylin and eosin to assess leukocyte infiltration. BALF was centrifuged and cell free supernatant stored at -80 °C for cytokine analysis by triplicate ELISA (R&D Systems). The cell pellet was resuspended in PBS and cells were counted using a Coulter counter or inspected by light microscopy.

### *A. thaliana* analyses

Arabidopsis (*Arabidopsis thaliana* Col-0) plants or seedlings were grown as described (Brock et al, 2010). ROS measurements and qPCR were performed as described (Albert & Furst, 2017; Brock et al, 2010) in 1 mm x 4 mm leaf pieces of 5-week-old Arabidopsis plants floated overnight in water, placed in a 96-well-plate (two pieces/well) containing 90 μl of the reaction mix (20 μM LuminolL-012, Wako Chemicals USA, 5 μg/ml horseradish peroxidase, Applichem, Germany) stimulated and the measured.. Further details are described in Supplemental Information.

### *In silico* docking and molecular modeling of TLR2-chitin interactions

The human TLR2 receptor PDB entry 2z7x (Jin et al, 2007) was used to generate input files for AutoDOCK 3.05 (126x64x64 grid points, resolution = 0.375 Å) and Autodock Vina (48x24x24 Å) (Trott & Olson, 2010) using AutoDockTools (Morris et al, 2009). MD simulations were performed in explicit solvent for 50 ns at 310 K using YASARA (Krieger & Vriend, 2015). The MD trajectories were analyzed using Conformational Analysis Tools (www.md-simulations.de/CAT/). The TLR2-SSL3-inhibited structure corresponds to PDB entry 5d3i (Koymans et al, 2015). Structures were inspected and figures and movies generated using VMD and Pymol 1.4.1. (Schrödinger). Further details are given in Supplemental Information.

### Statistical analysis

Experimental data was analyzed using Excel 2010 (Microsoft) and/or GraphPad Prism 6 or 7 (GraphPad Software, Inc.), microscopy data with ImageJ and Fiji, flow cytometry data using FlowJo software version 10. p-values were determined (α=0.05, β=0.8) as indicated. Principal component analysis was conducted as described in Expanded View methods and Expanded View code. p-values < 0.05 were generally considered statistically significant and are denoted by * throughout even if calculated p-values were considerably lower than p=0.05.

## Acknowledgements

This work as supported by the Wilhelm Schuler Stiftung, the University of Tübingen Medical Faculty, the University of Tübingen Graduate College “Of Plants and Men", the DFG-funded CRC 685 “Immunotherapy” and CRC/TR 156 “The skin as a sensor and effector organ orchestrating local and systemic immune responses", and the Federal State Baden-Württemberg Program “Glycobiology” (all to A.W.), NIH grant HL126094 (to C.D.C). Funded in part by grant R01AR069502 (L.S.M.) from the US National Institutes of Health. We thank Magno Delmiro Garcia, Johanna Bödder, Elisa Rusch, Timo Manz, Jennifer Ewald, Stefan Stevanović, Paul Vogel, Patrick Müller, Markus Löffler, P. Anoop Chandran, Sascha Venturelli, Christoph Mayer, Simon Anders, Jürgen Berger, Hubert Kalbacher, Martin Schaller, Beate Pömmerl, David Voehringer, Luigina Romani, Louris Feitsma, Eric Huizinga, Uffe Holmskov and Luke O’Neill for technical assistance, provision of reagents, critical reading of the manuscript and/or other very helpful comments.

## Authorship contributions

KF, YCG, OOW, FH, LS, CD, CT, SD, TMD, ZB, AS, DH, MAS, KK, TS, MK, CG, NAS, MF, SLL performed experiments; KF, YCG, OOW, FH, LS, CD, SD, TMD, ZB, AS, MS, TS, MK, CT, NAS, CG, BM, SLL, AAG, CDC, AW analyzed data; DH and AW wrote the manuscript which all other authors commented on; AW conceived and coordinated the study.

## Conflict of Interest Disclosures

KF, CG and AW are listed as inventors on two German patents. MF is an employee of Biognos AB, Göteborg, Sweden, and DH is an employee of Hoffmann La Roche, Basel, Switzerland. None of the other authors declares a conflict of interest.

## Abbreviations

Bronchio-alveolar lavage fluid (BALF), Calcofluor white (CFW), Concanavalin A (ConA), C-type lectin receptor (CLR), enhanced green fluorescent protein (EGFP), enzyme-linked immunosorbent assay (ELISA), human embryonic kidney (HEK), interferon (IFN), IFN regulatory factor (IRF), immunoglobulin (Ig), interleukin (IL), knockout (KO), lipopolysaccharide (LPS), median fluorescence intensity (MFI), microbe-associated molecular pattern (MAMP), myeloid differentiation primary response 88 (MyD88), N-acetyl-glucosamine (NAG, GlcNAc), nuclear factor ‘kappa-light-chain-enhancer’ of activated B-cells (NF-κB), pattern recognition receptors (PRRs), peripheral blood mononuclear cells (PBMCs), polymorphonuclear neutrophil (PMN), single nucleotide polymorphism (SNP), staphylococcal superantigen-like protein 3 (SSL3), TNF-associated factor (TRAF), Toll-like receptor (TLR), Tumor necrosis factor (TNF).

## Tables

Only supplemental tables are submitted, see Supplement File.

## Expanded View Section

### Expanded View Methods

#### Reagents and antibodies

All chemicals were from Sigma unless otherwise stated. Polymeric crude chitin, C4, C5, C6, C7 chitin oligomers and C10-15 chitosan oligomers were from Isosep, Elicityl and Carbosynth, respectively. According to the suppliers these were derived from crab shell biomass, chemically hydrolyzed and HPLC-fractionated oligomers >95% purity as confirmed by us using HLPC and mass spectrometry. Other PRR agonists were from Invivogen except flagellin from *Salmonella typhimurium* (Imgenex) and CpG 2006 from TIB Molbiol. Polymyxin B was from ThermoFisher. Recombinant mTLR2-Fc (a fusion of the murine TLR2 ECD and human IgG1-Fc) and hTLR2 ECD protein (His-tagged, no Fc fusion) were from R&D Systems. A control IgG1-Fc was produced in house using standard methods of antibody production. SSL3 was produced as described before (Koymans et al, 2015). In brief, a construct encompassing residues 134–326 of *S. aureus* strain NCTC 8325 SSL3 with a non-cleavable N-terminal His6-tag was expressed in *Escherichia coli* Rosetta-gami(DE3)*pLysS*, isolated under non-denaturing conditions from a HiTrap chelating HP column using an imidazole gradient for elution. SSL3 was stored in PBS and confirmed to be highly pure (>95%) by SDS-PAGE. For cellular assays endotoxin was first removed using Pierce High Capacity Endotoxin Removal Spin Columns. Ficoll was from Biochrom or Millipore.

#### Quality control of chitin and chitosan acetylation

The purity of purified chitin and chitosan preparations was >95% as stated by the manufacturers. C10-15 chitin was generated from C10-15 chitosan (2000-3000 MW, equivalent to 10-15 subunits) using sodium bicarbonate and acetic anhydride acetylation (Bueter et al, 2011). Briefly, chitosan 10-15 was suspended in 1 M sodium bicarbonate and 97% acetic anhydride was added, the reaction was carried out for 20 min at RT, followed by 10 min at 100 °C. The resulting degree of acetylation were assessed directly (C4 through C7 in water) or upon trifluoracetic acid hydrolysis (for C10-15, 2 h at 100 °C) by ESI and MALDI mass spectrometry and found to be >90% for the batches used here. Prior to ESI analysis samples were mixed with 50% methanol, 1% formic acid and injected via NanoES spray capillaries into an LTQ Orbitrap XL mass spectrometer (ThermoFisher). The spray voltage was set to 1.2 kV and mass spectra were acquired at a resolution of R = 30,000. Mass signals in MS1 scans were manually annotated and the mass error calculated in Xcalibur (ThermoFisher). Signal intensities were assessed to determine relative abundances of oligosaccharides present. Chitin batches were only used for further study if >90% of NAGs were acetylated. All mass spectra were recorded by MALDI-TOF-MS with a Reflex IV (Bruker Daltonics, Bremen, Germany) in reflector mode. Positive ions were detected, all spectra are the sum of 50 shots and a peptide standard (Peptide Calibration Standard II, Bruker Daltonics) was used for external calibration. 2,5-dihydroxybenzoic acid (DHB, Bruker Daltonics) was used as matrix, dissolved in water/acetonitrile/trifluoroacetic acid (50/49.05/0.05) at a concentration of 10 mg/ml. Before the measurement the samples were centrifuged and diluted with water (1:10). An aliquot of 1 μL of the samples was mixed with 1 μL of the matrix, the solution was spotted onto the MALDI polished steel sample plate and air-dried at room temperature. Prior to analysis chitin oligomers were suspended in endotoxin free water, tested for endotoxin contamination by using the limulus amoebocyte lysate (LAL) assay (Lonza, CH). Levels below 0.25 EU/ml (<25 pg/ml LPS) in final dilutions were considered ‘endotoxin-free’ and acceptable in comparison to other published studies (*cf.* Expanded View table S1). Additionally, stimulation of MoMacs or BMDMs with the equivalent 25 pg/ml LPS did not elicit any cytokine production upon overnight incubation. For levels >0.25 EU/ml the chitin preparation was incubated for 3 h with 2 mg/ml Polymyxin B (ThermoFisher), washed by centrifugation and re-assessed. In cellular assays Polymyxin B was used at 10 μg/ml. Fully de-acetylated chitosan 4,5,6,7 and 10-15-mers were inactive in initial experiments in THP-1 cells, primary MoMacs and BMDMs and thus not investigated further.

#### Chitin and zymosan depletion and Alexa647-coupling

C10-15 oligomers were also fluorescence-labeled using an aldehyde-reactive Alexa647 hydrazide cross-linker to react with the reducing end of the chitin chain (ThermoFisher, A20502) adapting a previously published protocol (Imai et al, 2002). For chitin and zymosan “depletion", 500 μg of insoluble fraction of zymosan or 200 μl of 1 mM C10-15 were resuspended in 2 ml 10 M NaOH and boiled at 95°C for 30 min as described (Gantner et al, 2003). Upon hot-alkali treatment zymosan and chitin were washed with 3 times with PBS or until pH was neutral. Subsequently the preparations were resuspended in endotoxin-free water (Braun). Chitinase treatment was performed using 30 μl 1 mM C10-15 or 100 μg/ml zymosan in water adding the volatile pyridine/glacial acetic acid buffer (pH 6.5) to a final concentration of 20 mM per 50μl reaction. 0.25 units chitinase from *Streptomyces griseus* (Sigma)., an exo-chitinase releasing di-NAG units from longer chitin chains, were added to zymosan and incubated for 18 h at 37°C. For MALDI analysis preparations were centrifuged and supernatants containing highly soluble di-NAGs were measured upon evaporation of the buffer.

#### Plasmid constructs

Plasmids for the expression of Flag-tagged TLR2 and TLR9 were gifts from I. Bekeredjian-Ding and A. Dalpke (Medical Microbiology, Heidelberg University, Germany). Plasmids expressing human NOD-2 were from T. Kufer (Hohenheim University, Germany) or for human Dectin-1 from G. Brown (Aberdeen University, UK). Plasmids were propagated and prepared according to standard procedures (Promega PureYield Midiprep). TLR2 hydrophobic pocket mutants were generated using the QuikChange II XL site-directed mutagenesis kit (Agilent) according to the manufacturer’s instructions using primers shown in Expanded View table S4. The correct insertion of the desired mutation as well as the absence of additional mutations in the coding region were confirmed by automated DNA sequencing (GATC Biotech).

#### Growth and fixation of *Candida albicans*

*C. albicans* strain SC5314 was stored as frozen stocks in 35% glycerol at -80°C and routinely grown on YPD agar (1% yeast extract, 2% bacteriological peptone, 2% glucose, and 1.5% agar) plates at 25°C. To prepare cultures, a single colony was inoculated in synthetic complete media (Formedium SC broth 2% glucose, additionally supplemented with 25 mg/L adenine sulfate) and shaken at 150 rpm at 30°C overnight. For fixation paraformaldehyde was added to a final concentration of 3.7% directly to the culture for an additional hour. Cells were harvested by centrifugation, washed twice in sterile Dulbecco’s phosphate-buffered saline (DPBS) and counted.

## Analysis of TLR2 binding *Candida albicans* cells

1x106 fixed *C. albicans* cells were incubated with 1 μM recombinant mouse TLR2 Fc chimera protein (R&D 1530-TR-050) or a corresponding control human IgG1 Fc (produced in house) while rocking overnight. On the next day, cells were stained with Alexa Fluor 594-conjugated mouse anti-human IgG (Jackson Immunoresearch 209-585-098, 1:10 dilution, 2 h incubation), 25 μM Calcofluor White and 50 μg/mL Concanavalin A CF488A (Biotum 20967 and 29016, 30 min incubation). All incubation steps were performed at room temperature in 100 μl DPBS in the dark. Washing was performed twice after each fluorophore incubation step with 200 μl DPBS by pelleting cells at 1000xg for 5 min. For microscopy, 200x103 cells were attached to Poly-L-Lysine coated coverslips (Corning 354085) in ProLong mount (Thermo Fisher P36965). Z-stacks were acquired with 0.2 μm step size on a Nikon Eclipse Ti2 system at 100x magnification with NIS-Elements 5.0 software and subjected to deconvolution. Images of individual cells were processed with the Fiji package for ImageJ (identical scaling of brightness/contrast, projection of maximum intensity of 15 selected stacks). Flow cytometry was conducted with the residual sample on a BD LSRFortessa^™^ system with 405, 488 and 532 nm laser excitation for respective fluorophores. Further settings upon request.

### Flow cytometric analysis of TLR2 chitin interaction

0.5 μM Alexa647-labeled C10-15 was incubated with 70 nM of recombinant mouse TLR2 Fc chimera protein (R&D 1530-TR-050) or a corresponding control IgG1-Fc in 200 μl DPBS over night at 4°C. Next day samples were stained with 1:15 diluted human Fcy-specific PE-conjugated F(ab’)2 fragments (Jackson Immunoresearch 109-116-098) in the dark at 4°C for 2 h. After washing twice with 1 ml DPBS (C10-15 precipitates were pelleted for 10 min at maximum speed) samples were measured on a BD FACSCanto^™^ II system with 488 and 633 nm laser excitation for the respective fluorophores. Further settings upon request.

### Microscale thermophoresis analysis of TLR2 chitin interaction

Microscale thermophoresis has been described elsewhere (Jerabek-Willemsen et al, 2011) and was conducted with fluorescence-labeled chitin and recombinant TLR2. In brief, 1:1000 diluted Alexa647-labelled C10-15 chitin (c = 17.7 μM) was mixed 1:1 with 2-fold serial dilutions of mTLR2-Fc or hTLR2 (both R&D Systems, c = 1150 nM (=417 μg/ml) or 1900 nM (=125 μg/ml) maximal concentrations in PBS converted to molar concentrations by using MW provided by R&D: 180.8 kDa and 67.5 kDa), incubated in the dark at RT overnight with gentle shaking. Solutions were then transferred to normal, non-coated Nanotemper capillaries and measured using 20%, 40% and 80% laser power on a Monolith NT.115 instrument and dataset with the highest ΔF fitted using Nanotemper software. Data were plotted with GraphPad Prism.

### Dual NF-kB luciferase assay and immunoblot in HEK293T cells

HEK293T were cultured in DMEM supplemented with 10% fetal calf serum, L-glutamine (2 mM), penicillin (100 U/ml), streptomycin (100 μg/ml) (all from ThemoFisher Scientific) at 37 °C and 5% CO2. 75.000 HEK293T cells were plated (24-well format, Greiner Bio One) and transiently transfected the next day with 25 ng TLR2 plasmid, 10 ng TLR9 plasmid, 0.25 ng NOD-2 plasmid or 50 ng Dectin-1 plasmid or the same amount of empty vector (EV) backbone, together with plasmids for firefly luciferase under the NF-κB promoter (100 ng) and constitutive Renilla luciferase reporter (10 ng). An EGFP expressing plasmid (100 ng) was used to monitor transfection efficiency which was usually >90%. Each set of transfection was adjusted with empty vector to equal amount of total plasmid. 32 hours post transfection, cells were stimulated for 18 h with the respective PRR agonists (MDP 200 nM, zymosan 100 μg/ml, CpG, Pam2 or 3 and C10-15 at 1 μM). Cells were washed in PBS, and subsequently lysed in 60 μl passive lysis buffer (Promega). Lysates were harvested by centrifugation and 10 μl of lysate used to measure luciferase activity on a Fluostar luminescence plate-reader (BMG Labtech). For immunoblot, 100 ng of TLR2 plasmid were transfected instead, and the cells lysed 48 h later in RIPA buffer (20 mM Tris Base, 150 mM NaCl, 1 mM EDTA, 1% (v/v) Triton, 10% (v/v) glycerol, 0.1% (w/v) SDS). Following BCA assays, equal amount of total lysate (20 μg) were loaded on 8% Tris-Glycine gels, separated, blotted on nitrocellulose membrane and probed with anti-Flag (Sigma F7425) and anti-rabbit IgG-HRP (Vector laboratoroies PI-1000). Blots were washed and incubated on a BlotCycler (Biozym) and bands detected using ECL reagents (Peqlab) on a Vilbert Lourmat CCD system.

### Respiratory burst (ROS) assay and cytokine analysis from Dectin-1-deficient immortalized macrophages

Immortalized murine macrophage progenitors (MOPs) from *CLEC7A* KO (Dectin-1 deficient) and the respective WT mice were originally described in (Rosas et al, 2008) and were a kind gift of P. Taylor (Cardiff University, UK). MOPs were cultured in complete RPMI 1640 with 1 μM β-estradiol (Sigma Aldrich), 10 ng/ml GM-CSF (Peprotech) at 37 °C and 5% CO2. Cells were differentiated using 20 ng/ml M-CSF (Peprotech) into immortalized macrophages (iMacs) (Rosas et al, 2008). For ROS assays, 40.000 cells of M-CSF iMacs were plated in 200μl into black 96-well plates with clear tissue-culture treated bottoms (Biozym). Subsequently, the cells were treated with PRR agonists and DCF added to 10 μM. ROS production was then quantified in triplicate and real time using a Fluorstar Optima Plate reader (BMG Labtech) set to 20 flashes per measurement (every 5 min), gain 1000 (top optic), excitation at 485 nm, emission at 535 nm. Counts for the unstimulated control were subtracted. For ELISA 40.000 M-CSF-differentiated iMacs were plated in 200μl in standard 96-well plates (Greiner Bio-One), treated with PRR agonists (C10-15 and Pam2 at 5 μM, zymosan at 50 μg/ml and PMA at 1 μg/ml) and supernatants collected for ELISA.

### Dimethyl-labeling of THP-1 cells and global phosho-proteome mass spectrometry analysis

THP-1 cells (Invivogen) were grown in complete RPMI medium and differentiated with 300 ng/ml PMA for overnight. The next day the medium was exchanged to PMA free complete media and the cells rested for another 48 h before stimulation with PBS, 5 μM C10-15 or 5 μM Pam2 for 30 minutes. After washing with ice-cold PBS (containing phosphatase and protease inhibitors, Roche), cells were lysed in 800 μl of lysis buffer (6 M guanidinium hydrochloride, 5 mM Tris(2-carboxymethyl)phosphine, 10 mM chloroacetamide, 100 mM Tris-HCl pH 8.5) heated to 99°C for lysis, reduction and alkylation of proteins. Cells were incubated for 10 minutes at 99°C, sonicated for 10 minutes and proteins digested with Trypsin overnight at room temperature. Protein concentration was determined using a Bradford assay. 7.9 mg total lysate were desalted on Sep-Pak C18 cartridges and dimethyl-labeled as published (Boersema et al, 2009). Prior mixing of protein samples, label incorporation and proper mixing was checked. The sample was fractionated on an offline Ultimate 3000 liquid chromatography system equipped with xBridge BEH C18 130A, 3.5 μm, 4.6 x 250 mm column operated under high pH using 5 mM NH4OH buffers. 60 fractions were collected and each fraction was analyzed by LC-MS/MS for full proteome analysis. The 60 fractions were concatenated into 15 pools and dried by vacuum centrifugation. Peptide pools were reconstituted in 1 ml of 80% ACN, 6% TFA and enriched with TiO2 spheres for 30 minutes in a protein to bead ratio of 2:1. Pelleted beads were washed with 100μl of 30% ACN, 1% TFA followed by 80% ACN, 1%TFA. Elution from the beads was performed two times with 100 μl of 5% ammonia hydroxide solution in 60% acetonitrile (pH > 10.5) and once with 10 μl 1% formic acid in 80% ACN. Acetonitrile was eliminated by vacuum centrifugation and phospho-peptides desalted on Stagetips prior LC-MS/MS measurements. LC-MS/MS analyses were performed on an Easy-nLC 1200 equipped with a 20 cm and 75 μm ID PicoTip fused silica emitter packed with ReproSil-Pur C18-AQ 1.9 μm resin (Dr. Maisch Ltd.) coupled to an Q Exactive HF (both Thermo Scientific). Phospho-peptide enriched samples were separated using a 87 minute segmented gradient of 5-50% solvent B (80% ACN in 0.1% formic acid) at a constant flow rate of 200 nl/min while high pH LC fractions were separated by an 35 minute gradient. The Q Exactive HF was operated in positive mode. Full scans were acquired from m/z 300 to 1,650 with a resolution of 60,000. For proteome measurements the 20 most intense ions and for phospho-proteome the seven most intense ions were fragmented by higher energy collisional dissociation (HCD) and MS/MS spectra recorded with a resolution of 30,000 and 60,000, respectively. The MS data were processed using default parameters of the MaxQuant software suite (v1.5.5.1) (Cox & Mann, 2008). Extracted peak lists were submitted to database search using the Andromeda search engine (Cox et al, 2011) to query a target-decoy database of *H. sapiens* proteome (release 2014_02; 88,692 entries). In database search, full tryptic specificity was required and up to two missed cleavages were allowed. Carbamidomethylation of cysteine was set as fixed modification; protein N-terminal acetylation, oxidation of methionine, and phosphorylation of serine, threonine, and tyrosine were set as variable modifications. Overall light-, medium- and heavy-demethylation labeling on lysine residues and peptide N-termini were defined. “Re-quantify", calculating ratios for isotope-patterns not assembled into labeling triplets, was enabled. Initial precursor mass tolerance was set to 4.5 ppm at the precursor ion and 20 ppm at the fragment ion level. False discovery rates were set to 1% at peptide, phosphorylation site, and protein group level. The MaxQuant output was analyzed using the Perseus software (1.5.3.2). Pearson correlation coefficient was calculated for replicates. Differentially expressed proteins and phosphorylation events were identified using the two-tailed, signal intensity-weighted “Significance B” test (p ≤ 0.01). Proteome and phospho data were mapped to pathways using the KEGG Pathway Mapper (http://www.genome.jp/kegg/tool/map_pathway2.html) and manually annotated regarding the regulation of individual proteins or phospho-sites. The mass spectrometry proteomics data have been deposited to the ProteomeXchange Consortium via the PRIDE partner repository at www.ebi.ac.uk/pride with the dataset identifier PXD007542.

### THP-1 cell line analysis

The human monocytic THP-1 cell line (Invivogen) and two THP-1-based TLR2 Cas9-CRISPR-edited clones or their parental cell lines (a gift from V. Hornung, Gene Center, Munich, Germany, (Schmid-Burgk et al, 2014)) were maintained at 37 °C and 5% CO2 and differentiated into macrophages by exposure to 300 ng/ml PMA in complete RPMI overnight and then rested for an additional 48 hours in PMA-free complete RPMI medium. For stimulation C7, C10-15, Pam2, Pam3 were used at 5 μM, LPS at 1 μg/ml, zymosan preparations at 25 μg/ml. For competition experiments, C10-15 was used at 5 μM and C5 at 0.2, 1 and 5 μM and pre-incubated for 2 h. For qPCR analysis, 1x106 cells were seeded and stimulated in 24 well plates as indicated, washed with PBS, harvested in RLT buffer and total mRNA isolated according to the RNeasy kit (Qiagen) on a Qiacube robot. Upon reverse transcription to cDNA (High Capacity RNA-to-cDNA Kit; ThermoFisher), mRNA expression of the indicated cytokines and chemokines was quantified in triplicates relative to TBP using gene-specific TaqMan primers (ThermoFisher, Expanded View table S5) on a real-time cycler (Thermo QuantStudio 7 Flex). For ELISA, 50,000 cells per 96-well triplicate (Greiner Bio-One) or 500,000 in 24-well plates (both Greiner Bio-One) were seeded and stimulated overnight. Subsequently, IL-6 and TNF ELISAs were performed in triplicates (Biolegend).

### Human study subjects and sample acquisition

All healthy blood donors included in the analyses of immune cells for this study provided their written informed consent before study inclusion. Approval for use of their biomaterials was obtained by the local ethics committee at the University of Tübingen, in accordance with the principles laid down in the Declaration of Helsinki. Buffy coats were obtained from blood donations of healthy donors and received from the Center for Clinical Transfusion Medicine (ZKT) at the University Hospital Tübingen and whole blood from voluntary healthy donors was obtained at the University of Tübingen, Department of Immunology.

### Mice

*Myd88* and *Tlr2 KO* mice (both on a C57BL/6 background, a gift from H. Wagner, Ludwigs-Maximilian University, Munich) and C3H/HEJ (*Tlr4*^*LpsD*^, Jackson) and their respective WT counterparts (WT C57BL/6 or C3H/HEN, respectively) were used between 8 and 20 weeks of age in accordance with local institutional guidelines on animal experiments and under specific locally approved protocols for sacrificing and *in vivo* work. All mouse colonies were maintained in line with local regulatory guidelines and hygiene monitoring.

### Isolation of PBMCs from human peripheral blood

Peripheral blood mononuclear cells (PBMCs) were isolated from heparinized whole blood (Department of Immunology) or buffy coats (University Hospital Tübingen Transfusion Medicine) from healthy human volunteers by standard density gradient separation (Biocoll, Biochrom GmbH, Berlin, Germany), washed twice in PBS and counted with trypan blue.

### Isolation and analysis of primary human monocyte-derived macrophages

Primary monocytes were isolated from PBMC using standard Ficoll density gradient purification and washed three times with RPMI. For macrophage differentiation, monocytes were purified by positive selection from PBMC using anti-CD14 magnetic beads (Miltenyi Biotec, >90% purity assessed by anti-CD14-PE flow cytometry) and seeded in RPMI-1640 (Sigma), 10% FCS (GE Healthcare), 2 mM L-Glutamine (ThermoFisher), 1 % Pen Strep (ThermoFisher) at a concentration of 0.75x106 cells/ml in 96-well (ELISA) and/or 24-well (electron microscopy) tissue culture plates. Cells were differentiated into macrophages in the presence of 25 ng/ml recombinant human GM-CSF (Prepro Tech) for 6 days (Verreck et al., 2004) and seeded and treated as described. Stimuli were as follows: LPS at 100 ng/ml, C10-15, Pam2 and Pam3 all 5 μM, R848 at 5 μg/ml. For competition experiments, C10-15 was used at 5 μM and C5 at 0.2 μM pre-incubated for 2 h. For electron microscopy (performed by J. Berger, Max-Planck-Institute, Tübingen, Germany) cells were seeded on poly-L-lysine-treated cover slips in 24 well plates. Specimen preparation and EM analysis was done according to standard procedures. For RNAi experiments, all siRNAs (see Expanded View table S6) were from Dharmacon (siGENOME, GE Dharmacon) and resuspended in RNase/DNase-free H2O to a stock concentration of 10 μM according to manufacturer’s instructions. Aliquots were stored at -20 °C until use. On day 5 of GM-CSF differentiation, the medium was replaced and differentiated MoMacs were transfected with 35 nM siRNA using Viromer Blue (Biozol) according to manufacturer’s instructions for adherent cells. Transfections were carried out in triplicate wells. On day 6, cells were stimulated as mentioned before and supernatants harvested on day 7 and analyzed by ELISA.

### Isolation and analysis of primary human neutrophils

Primary neutrophils were isolated from the neutrophil-erythrocyte pellet after Ficoll density gradient purification. Erythrocytes were lysed twice using 1x Ammonium chloride buffer (10X stock: pH 7.3, 1.54 M NH4Cl, 100 mM KHCO3, 1 mM EDTA) first for 20 min and after subsequent centrifugation for another 10 min at 4 °C. The cells were resuspended in complete RPMI and 1x106 cells/well were seeded (24-well plate). Purities were typically >97% and not pre-activated as assessed by PE anti-CD15, FITC anti-CD66b and BV421 anti-CD62L (all Biolegend) flow cytometry and appropriate isotype controls. After resting at 37°C and 5% CO2 for 30 min, the PMNs were stimulated with PRR agonists C10-15 (5 μM), PMA (500 ng/ml), LPS (200 ng/ml), Pam2 and Pam3 (0.5 or 5 μM), zymosan and zymosan depleted (both 25 μg/ml) for 1 h for CD62L shedding analysis using anti-CD62L-BV421 Abs. For blocking experiments, the cells were pre-treated with a purified anti-mouse/human CD282 (TLR2) Ab (Biolegend 121802, (Meng et al, 2003)) or an anti-HA (H9658) isotype control (Sigma), both at 10 μg/ml, or with SSL3 (5 nM) for 30 min before PRR ligand addition. Thereafter FACS staining using BV421 anti-CD62L and an isotype control was performed and the samples were then analyzed with a BD FACSCanto II. For ROS assays, 2x105 cells per 96-well were seeded in 100 μl and 100 μl DCF at 10 μM together with different stimuli at 2x concentration added. Fluorescence was measured continuously for 3 h but for clarity only final fluorescence counts were plotted. For ELISA, purified PMNs were seeded at 1x106 per 24-well in complete RPMI, stimulated for 4 h and supernatants were harvested for IL-8 ELISA (Biolegend).

### Human whole blood microarray and qPCR analyses

For whole blood assays 1 ml freshly drawn heparinized blood was aliquoted into 24 well plates and stimulated for 3 h. Subsequently, plasma was removed and erythrocyte lysis performed. The remaining cells were frozen in RLT at -80 °C for further q-PCR analysis according to the QIAamp RNA Blood Mini Kit. mRNA was isolated using the same kit on a Qiacube robot, genomic DNA digested (Turbo DNA-free Kit, ThermoFisher), transcribed to cDNA (High Capacity RNA-to-cDNA Kit; ThermoFisher). For microarray analysis (performed by A. Witten, Münster University, Germany) total RNA and amplified RNA were quantified on a NanoDrop 8000 spectrometer (Peqlab) and quality controlled using the Nano 6000 Kit for Bioanalyzer (Agilent). aRNA synthesis was conducted using the TargetAmp-Nano Labeling Kit for Illumina Expression BeadChip (epicentre) and analysis carried out using the Illumina HumanHT-12 v4.0 Expression BeadChip Kit on an Illumina HiScan array scanner. Microarray data was analyzed using R version 3.4.2. The supplementary PDF holds all R code used to produce the respective figures, plus additional illustrations on how we interpret these data. Briefly, the analysis resulting in Fig. 3 is as follows. Microarray data was processed with R package ‘limma’ version 3.34.8. To compute PCA, 6,191 variable genes were selected with limma’s F-test (ANOVA) and their normalized, log2-transformed expression values used as input for R’s prcomp function. Differentially expressed genes were computed between chitin and other treatments using limma at an FDR < 0.05 after Benjamini-Hochberg correction. From these, the Heatmap in Fig. 3 displays genes with a fold change of 1.1 (Chitin > all) and 1.2 (Chitin up and/or down in PAM2 and PAM3). The microarray data have been deposited to the Gene Expression Omnibus (GEO) repository at www.ncbi.nlm.nih.gov/geo/ with the dataset identifier GSE103094. For qPCR, mRNA expression of several cytokines and chemokines was quantified in triplicates relative to TBP using gene-specific TaqMan primers (ThermoFisher, Expanded View table S5) on a real-time cycler (Thermo QuantStudio 7 Flex).

### Isolation and analysis of mouse bone marrow derived macrophages (BMDM)

Mice were sacrificed and femurs and tibia excised. The bones were then opened and bone marrow (BM) cells were flushed into a 50 mL Falcon tube with RPMI+10% FCS medium with a 5 mL syringe and a 27 g needle. The cells were then passed through a 0.22 μM strainer, followed by centrifugation and counting of the cells. 6x106 BM cells/well were seeded in a 6 well plate in RPMI +10% murine GM-CSF + 10% FCS +1% Penicillin/Streptomycin + 1% L-Glutamine culture medium and incubated at 37°C and 5% CO2 Every second day fresh culture medium was added. After 6 days, the resulting BMDMs were detached from the plate using cold PBS with 0.5 mM EDTA, centrifuged and seeded in 96 well plates (0.75x106 cells/ml) for stimulation. The released TNF and/or IL-6 levels were measured after 18 h incubation with the following stimuli: C5 (5 μM), C6 (5 μM), C7 (5 μM), C10-15 (5 μM), LPS (1 μg/ml), poly(I:C) (20 μg/ml), Pam2 (5 μM), Pam3 (5 μM), zymosan and zymosan depleted (both 10 μg/ml). For measuring ROS activity 75,000 cells were seeded and assayed as described above for WT and *Clec7a* KO (Dectin-1 deficient) immortalized macrophages.

### *In vivo* dermal inflammation analysis

Female LysM-EGFP mice expressing EGFP in myeloid cells due to knock-in of EGFP into the lysozyme M gene were used as before described (Cho et al, 2012). The dorsal backs of anesthetized LysM-EGFP mice (2% isoflurane) were shaved and injected intradermally (i.d.) with chitin oligomers C10-15, C7 and C5 at 5 μM concentration in 100 μL of PBS, 1 mg/kg LPS in 100μL of PBS or 100 μL of PBS alone using a 29 gauge insulin syringe. For *in vivo* imaging of EGFP neutrophils appearing on the site of injection a Lumina III IVIS (PerkinElmer) was used. Mice were put into the imaging chamber of the system after being anesthetized using isoflurane. Images were acquired at time 0, 2, 6, hours and 1, 2, 3 days following injection. The EGFP-expressing neutrophils within the injection area were visualized using GFP filters for excitation (460 nm) and emission (520 nm) at an exposure time of 0.5 seconds. Analysis of the images was performed using Living Image 4.3 software (Perkin Elmer) and fluorescence intensities, expressed as average radiance (photons per cm^2^ per s) were measured by drawing circular region of interest over the entire site of injection. 10 mm punch biopsies were also taken from the injection site at 3 days after treatment. Skin samples were weighed then homogenized in a protein lysis buffer using mechanical homogenization. Homogenates were stored at -80 °C. Myeloperoxidase (MPO) ELISAs were performed as per manufacturer instructions using commercially available kits (R&D Systems).

### *In vivo* lung inflammation analysis

WT and *Tlr2* KO mice were anaesthetized using ketamine-xylazine solution and a vertical cut was made on the neck to expose trachea to instill 30 μl of 1 mM chitin oligomers using a Hamilton syringe. For competition experiments with SSL3, 25 μl of an endotoxin-free 50 μg/ml (2.6 mM) of SSL3 solution was first instilled into the trachea followed by instillation of 30 μl of a 1 mM C10-15 solution. The wound was afterwards sealed using Vetbond tissue adhesive. Mice were euthanized at 12 hours later to harvest BALF and lung tissues. Mice were euthanized 12 h post instillation of chitin to harvest BALF. Lung tissues were inflated at 1 atm pressure with 0.5% low melting point agar and fixed using formalin. Lung sections were embedded and sections were stained with hematoxylin and eosin to assess leukocyte infiltration. BALF was centrifuged and cell free supernatant stored at -80 °C for cytokine analysis by triplicate ELISA (R&D Systems). The cell pellet was resuspended in PBS and cells were counted using a Coulter counter or inspected by light microscopy.

### *A. thaliana* analyses

Arabidopsis (*Arabidopsis thaliana* Col-0) plants or seedlings were grown on soil or half-strength Murashige and Skoog, respectively (Brock et al, 2010). To measure the generation of reactive oxygen species in a chemiluminescent assay (Albert & Furst, 2017), 1mm x 4 mm leaf pieces of 5-week-old Arabidopsis plants were floated overnight in water and the next day placed in a 96-well-plate (two pieces/well) containing 90 μl of the reaction mix: 20 μM Luminol (L-012, Wako Chemicals USA), 5 μg/ml horseradish peroxidase (Applichem, Germany) and water. After equilibration the stimulus was added in a volume of 10 μl according to the concentration to be tested. Light emission was detected in a 96-well-plate reader (Mithras LB 940; Berthold Technologies) as relative light units. Light signals were measured every 1 sec per well and sample. To test inhibitory effects, C5 was added into the reaction mix in the concentration indicated. After 30 min C10-C15 was added. For measurement of transcript levels, RNA isolation and qRT-PCR analysis of plant material treated with different chitin fragments were performed as described (Brock et al, 2010) using gene specific primers or the defense marker gene *Flagellin-responsive kinase 1* (FRK1-100-F AGCGGTCAGATTTCAACAGT; FRK1-100-R AAGACTATAAACATCACTCT), *MLO12* (MLO-F ACGGTGGTTGTCGGTATAAGCC; MLO-R AGGGCAGCCAAAGATATGAGTCC) and *WRKY40* (WRKY40-F AGCTTCTGACACTACCCTCGTTG; WRKY40-R TTGACAGAACAGCTTGGAGCAC). *EF-1a* transcripts (EF1a-F TCACATCAACATTGTGGTCATTGG; EF1a-R TTGATCTGGTCAAGAGCCTACAG) served for normalization, and controls were set to 1. All quantifications were made in duplicate on RNA samples obtained from three independent experiments, each performed with a pool of 20-30 10 day old Arabidopsis seedlings.

### *In silico* docking and molecular modeling of TLR2 chitin interactions

The atom coordinates of human TLR2 receptor were extracted from PDB entry 2z7x (Jin et al, 2007). Chitin ligands were minimized using YASARA (Krieger & Vriend, 2015). AutoDockTools (Morris et al, 2009) was used for the preparation of the input files for AutoDOCK 3.05 and Autodock Vina (Trott & Olson, 2010). The dockings were performed to the hydrophobic cavity of TLR2. Box dimensions: AutoDOCK (126x64x64 grid points, resolution =0.375 Å), Vina (48x24x24 Å). For each ligand 300 dockings (Genetic algorithm/local search, 107 energy evaluations each) were performed using AutoDock 3.05 and all 300 poses obtained were analyzed. Dockings with Vina were performed with an exhaustiveness parameter of 100 and 20 poses were used as output for further analysis. Since the chitin ligands C4 and C5 initially did not give many poses inside the cavity, the dockings were repeated with F349 in the ‘gate open’ conformation (Expanded View figures S4e) which improved the outcome of the dockings significantly. The starting structures for Molecular Dynamics simulations of human TLR2 were prepared based on PDB entry 2z7x. The TLR2 protein was extracted from the PDB entry and solvated in 0.1% NaCl solution using the YASARA graphical interface (Krieger & Vriend, 2015) following standard procedures. The complexes with chitin C6, C7 and C10 were built based on the results obtained from the docking of C5. Additionally, MD simulations with C6 were performed with various orientations of the chitin chain in the binding site. MD trajectories were sampled for 50 ns at 310 K using the AMBER14 (Hornak et al, 2006) force field in YASARA, which includes Glycam-06 (Kirschner et al, 2008) parameters for carbohydrates. Snapshots were stored every 25 ps. The MD trajectories were analyzed using Conformational Analysis Tools (www.md-simulations.de/CAT/). The TLR2-SSL3 structure (Fig. 4f) corresponds to PDB entry 5d3i (Koymans et al, 2015). Structures were inspected and figures and movies generated using VMD and Pymol 1.4.1. (Schrödinger).

### Statistical analysis

Experimental data was analyzed using Excel 2010 (Microsoft) and/or GraphPad Prism 6 or 7 (GraphPad Software, Inc.), microscopy data with ImageJ and Fiji, flow data using FlowJo software version 10. p-values were determined (α=0.05, β=0.8) as indicated. p-values < 0.05 were generally considered statistically significant and are denoted by * throughout even if calculated p-values were considerably lower than p=0.05. PCA and microarray analyses are described in the above section on whole blood analyses and further detailed in the Expanded View section.

## Expanded View Figure Legends

**Expanded View figure S1.**
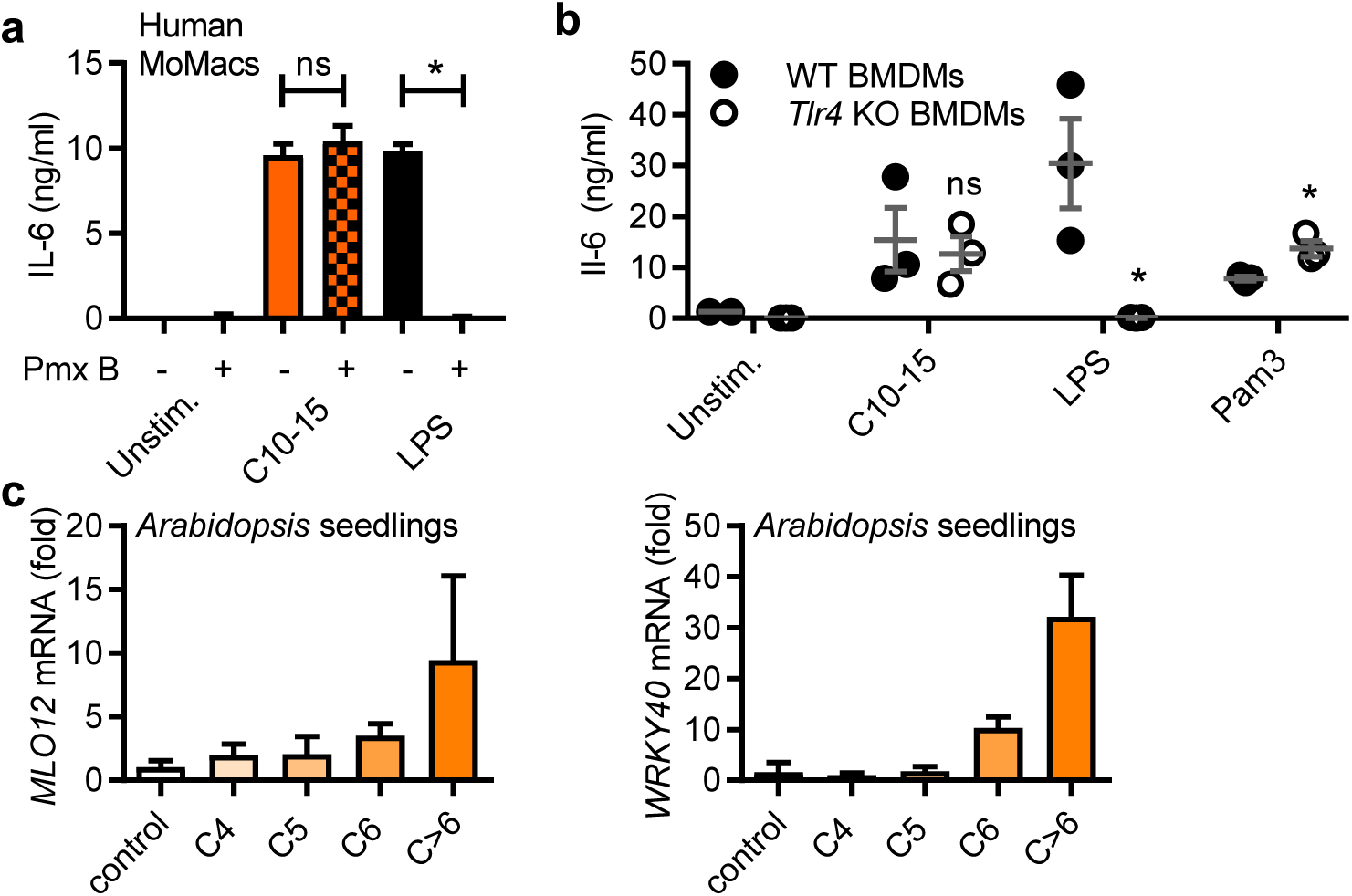
(a,b) Cytokine release induced by chitin is independent of endotoxin. (a) IL-6 release from stimulated human MoMacs stimulated in the absence or presence of polymyxin B (Pmx B, n=3), and (b) IL-6 release from WT or *Tlr4* KO BMDMs (n=3) are shown for one representative batch of C10-15. (c,d) Target gene induction in *A. thaliana* seedlings. *MLO12* (c) and *WRKY40* mRNA induction (n=3). b represents data (mean+SEM) combined from 3mice. In a and c one representative of ‘n’ (given in brackets for each panel) independent experiments is shown (mean+SD). * p<0.05 according to Student’s t-test (a,b).

**Expanded View figure S2.**
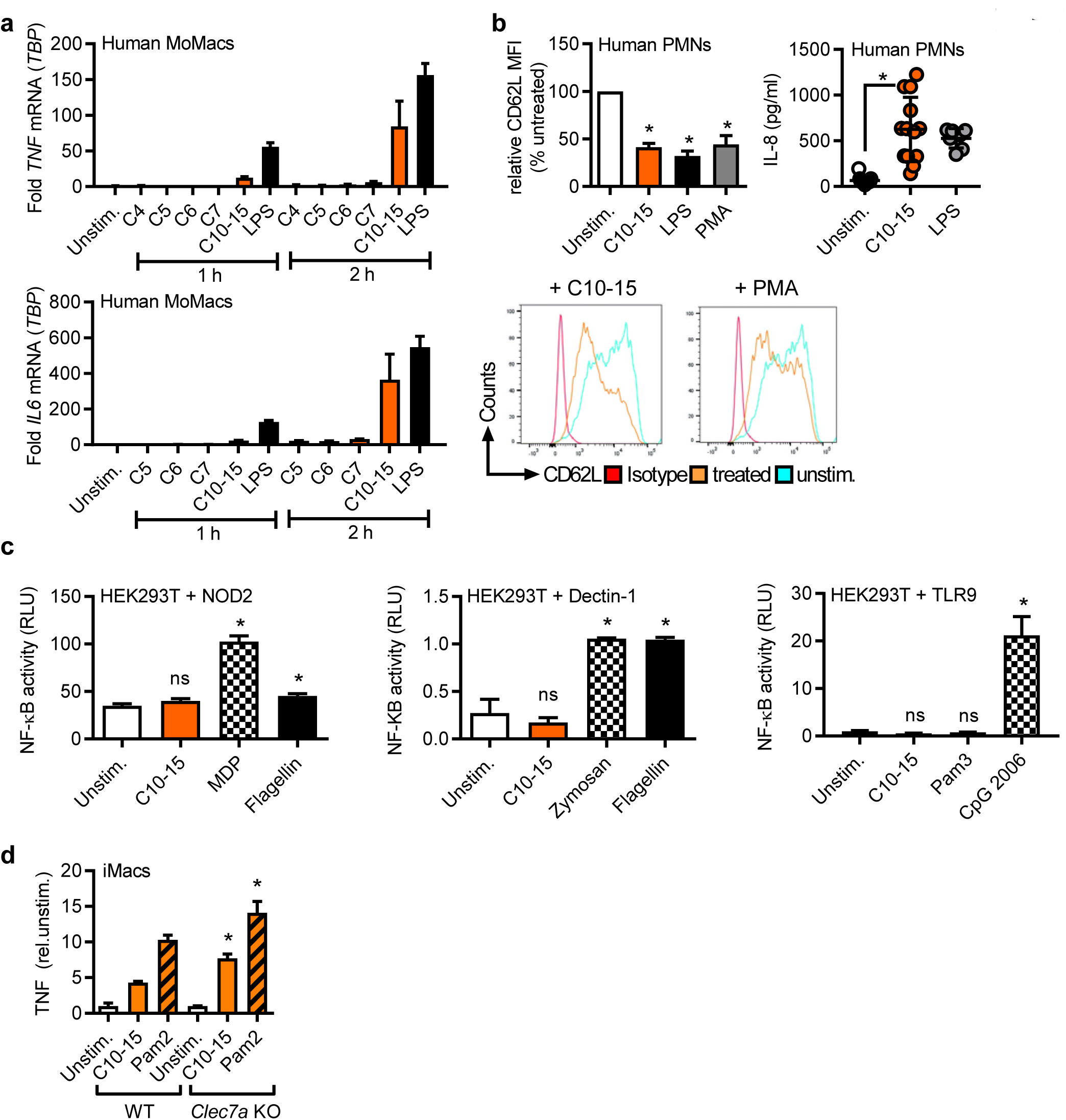
(a) *IL6* or *TNF* mRNA induction relative to *TBP* in human MoMacs (n=3) (b) Relative CD62L shedding (n=8, histograms shown for 1 representative donor) and IL-8 release (n=7-13) from human PMNs upon 1 or 4 h stimulation, respectively. (c) NF-κB activity in empty vector (EV) or NOD2, Dectin-1 or TLR9-transfected HEK293T cells (n=2-3). (d) TNF release (relative to unstimulated) from WT or *Clec7a* KO (Dectin-1 deficient) immortalized macrophages (iMacs, n=2). b represents data (mean+SEM) combined from 7-13 donors. In a, c and d one representative of ‘n’ (given in brackets for each panel) independent experiments is shown (mean+SD). * p<0.05 according to Wilcoxon signed rank sum (b) or Student’s t-test (c,d).

**Expanded View figure S3.**
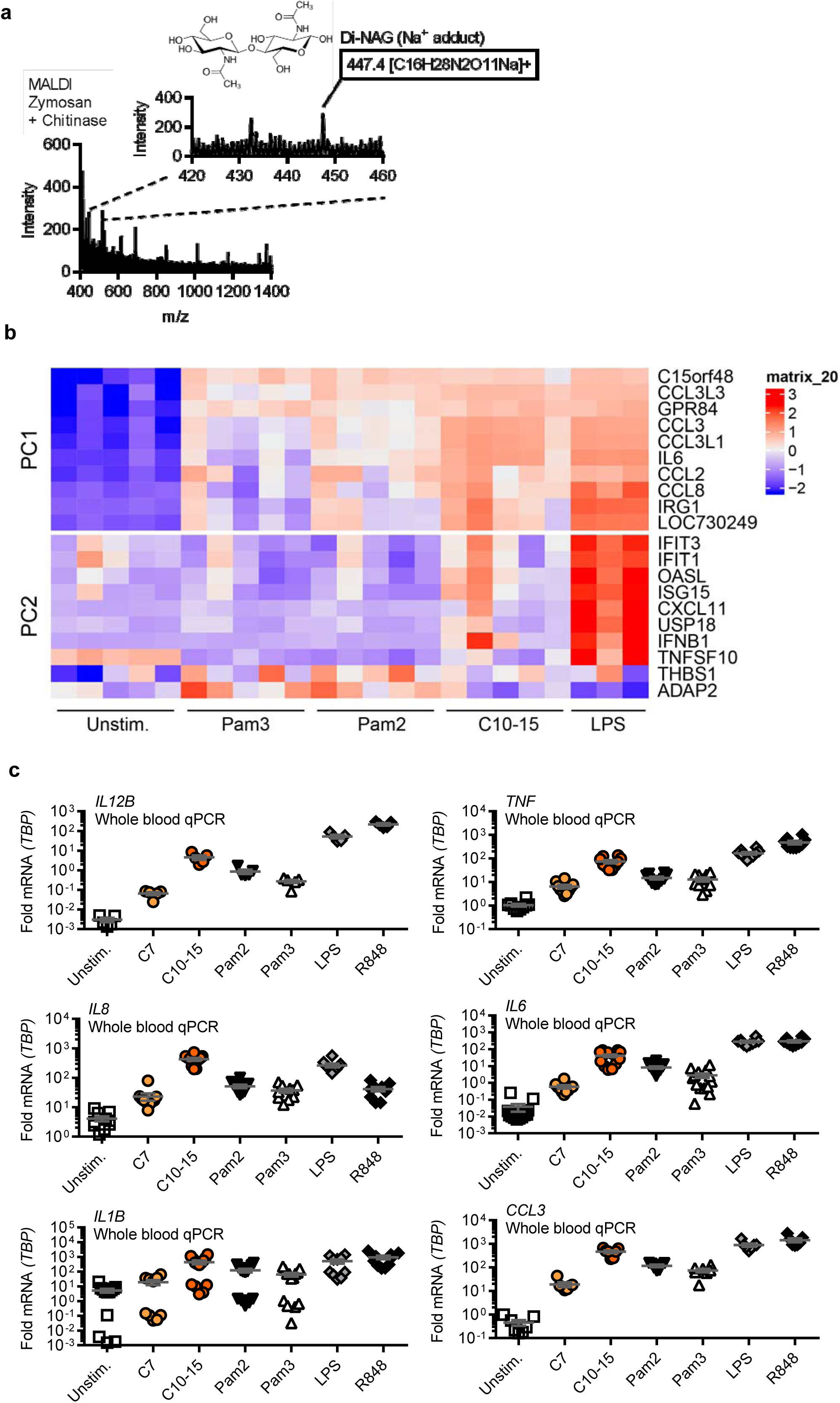
(a) MALDI-TOF-MS analysis of chitinase-treated zymosan supernatants (n=2). (b) Whole blood microarray (n=3-5 donors) intensities for selected genes contributing to PC1 or PC2. (c) Relative cytokine mRNAs in whole blood stimulation (n=5-13 donors). In a one representative of 2 independent experiments is shown (mean+SD). b represents data (mean+SEM) combined from 5-13 donors.

**Expanded View figure S4.**
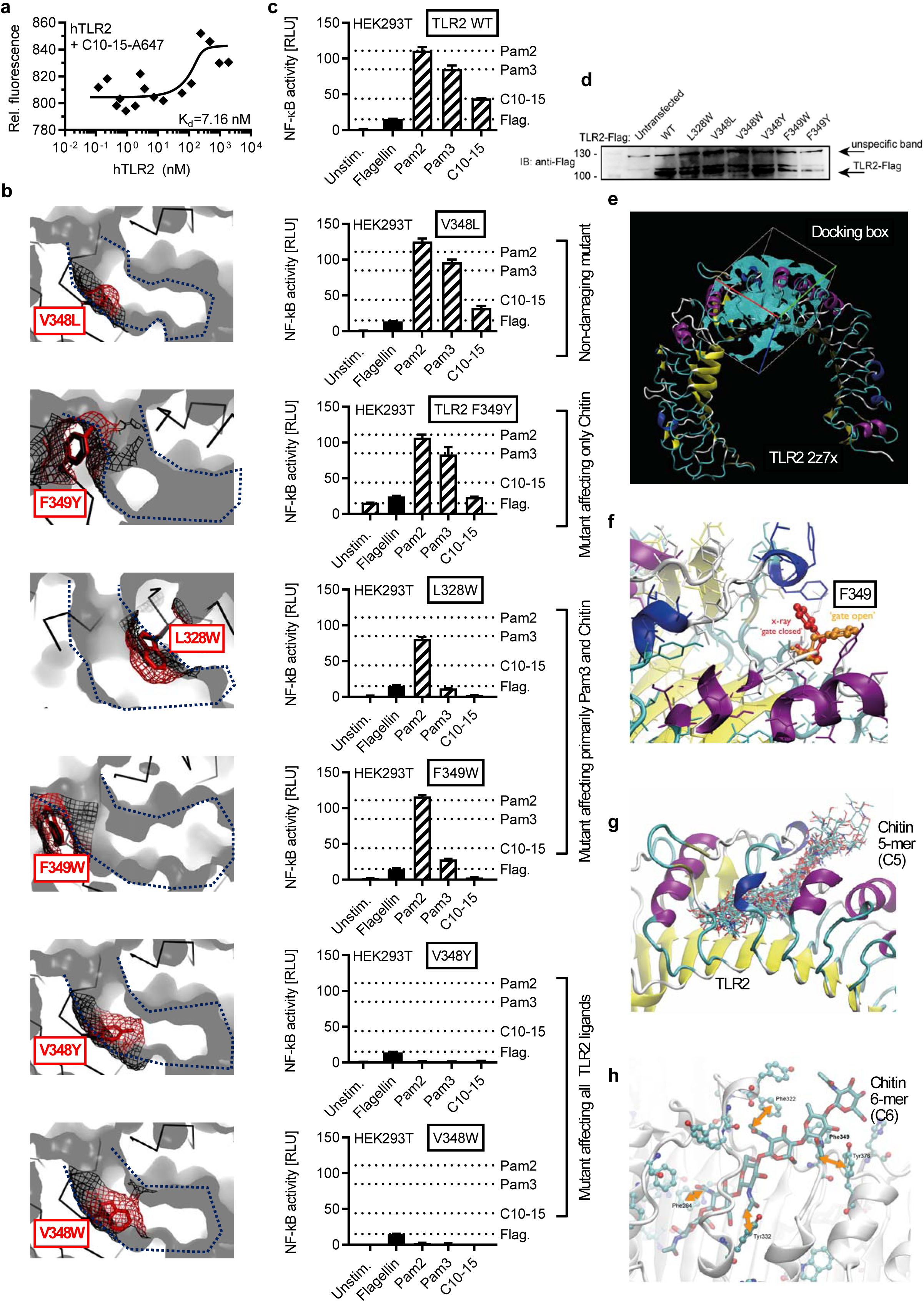
(a) Microscale thermophoresis analysis of Alexa647-labeled C10-15 and hTLR2 protein (n=3). Modeling (b), functional NF-κB activity (c) and expression levels (d) in empty vector (EV) or TLR2-transfected HEK293T cells (n=2) for TLR2 WT or point mutants designed to narrow the hydrophobic pocket. In b, a blue dotted line delineates the hydrophobic pocket, here shown in cross-section. Black mesh signifies original surface profile in pdb 2z7x, red mesh the predicted surface of the respective point mutant. In c dotted lines signify the levels of NF-κB activity observed for WT TLR2 for the respective ligand. (d) Expression of TLR2 mutants verified by anti-Flag immunoblot (n=1). (e) Docking grid used for placing of a chitin 5-mer. (f) Close-up on the gatekeeper F349 which restricts access to the cavity in ‘closed position’ (red) but allows access in ‘open’ rotamer position (orange). (g) Docking poses of a 5-mer (C5) in the grid specified in e and with F349 in ‘open’ rotamer position. (h) Potential interactions of chitin 6-mer (C6) hydrophobic methyl groups with aromatic side chains in the TLR2 hydrophobic cavity. In a, c and d one representative of ‘n’ (given in brackets for each panel) independent experiments is shown (mean+SD). * p<0.05 according to Student’s t-test (c).

**Expanded View figure S5.**
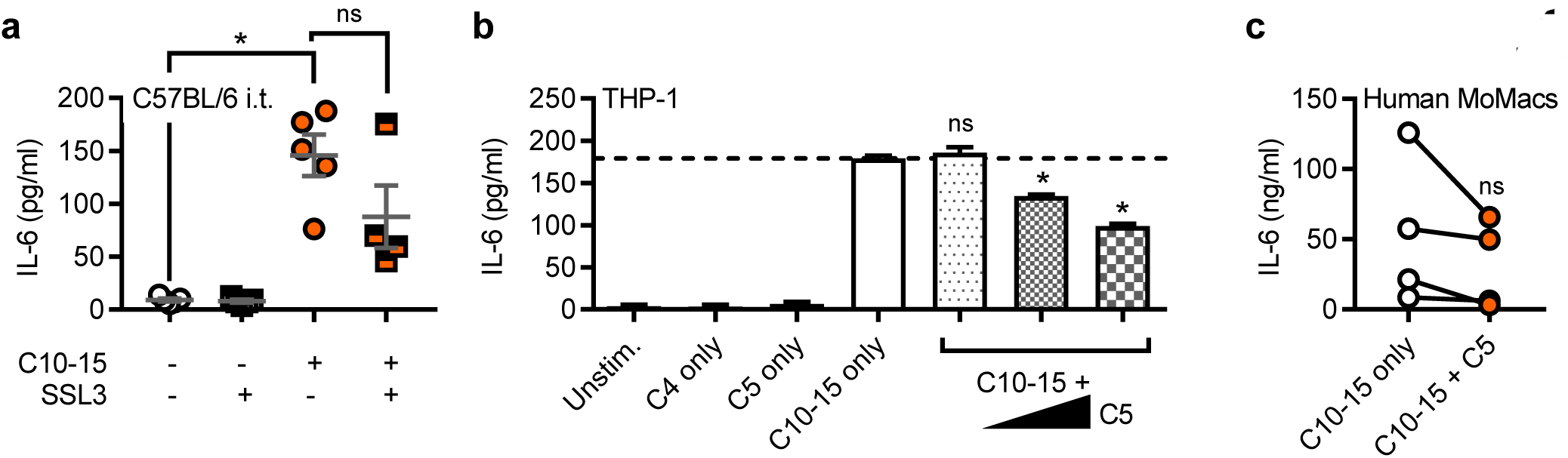
(a) BALF IL-6 in C57BL/6 mice (n=4-5/group) upon C10-15 administration without or with SSL3. IL-6 release from THP-1 cells (b) or primary MoMacs (c) without or with C5 (n=3 or 4). a and c represent data (mean+SEM) combined from ‘n’ biological replicates (human donors or mice, respectively). In b one representative of 3 independent experiments is shown (mean+SD). * p<0.05 according to one-way ANOVA with Dunnett’s multiple comparison (a), Student’s t-test (b) or Wilcoxon signed rank sum (c) test.

**Expanded View figure S6.**
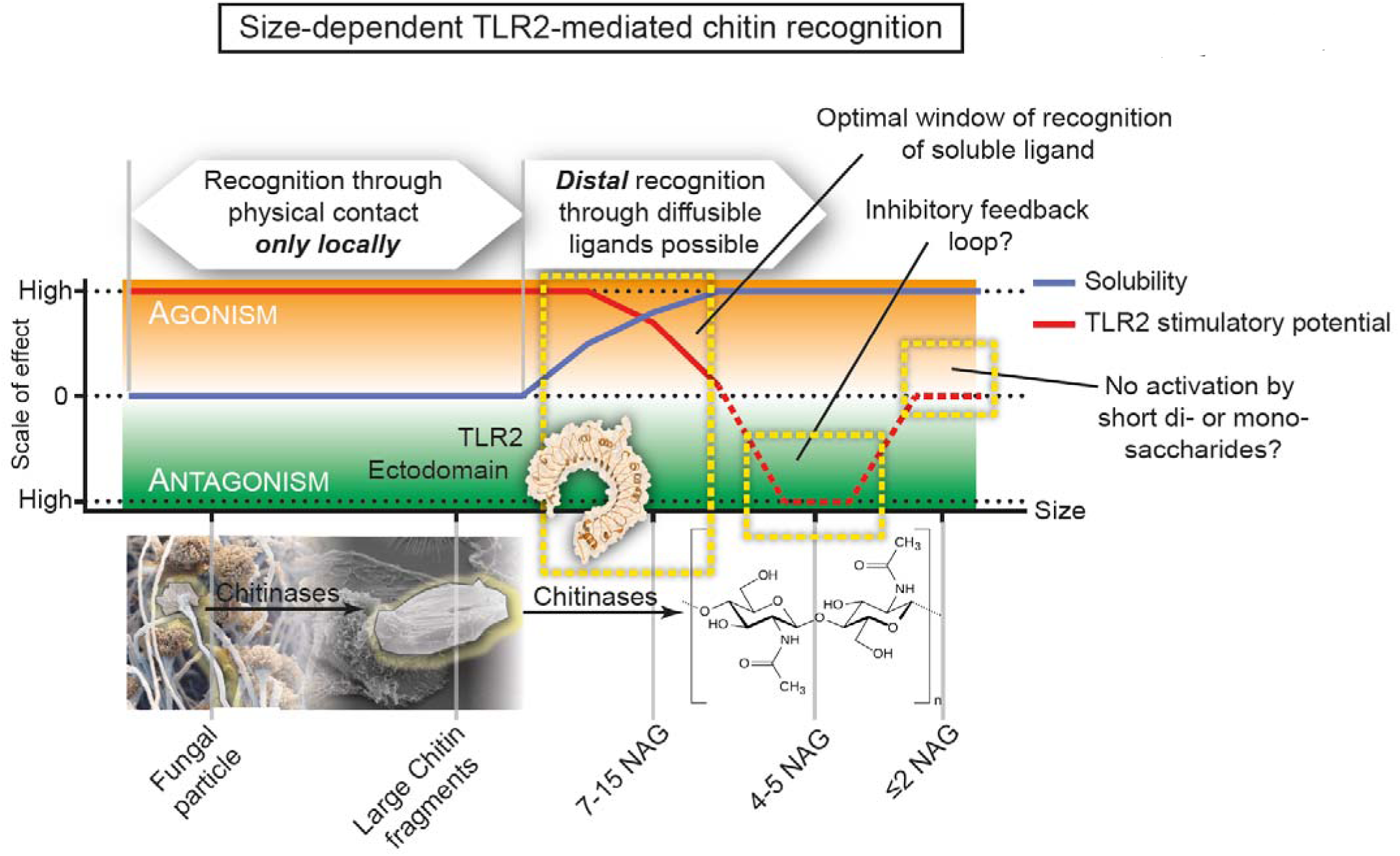
Schematic representation of size-dependent recognition of chitin oligomers by TLR2. Through the activity of chitinase or physical fragmentation a range of different sized fragments and oligomers is generated. TLR2 recognition is possible for both particulate chitin as well as diffusible oligomers. As recognition can occur in a window where both activatory potential and solubility are high, distal recognition of fungal pathogens is possible, whilst the minimum length required for agonism (6 NAG) ensures that mono- or disaccharides are prevented from receptor activation. As activating oligomers are cleaved, soluble fragments with antagonistic properties may arise, representing an inherent negative regulatory loop.

## Expanded View Movie Legends

**Expanded View movie S1: General concept of Pam3 and chitin binding to the TLR2 hydrophobic pocket** (uploaded as mp4 video file)

**Expanded View movie S2: Steered docking and molecular dynamics simulation of a chitin 7-mer into the TLR2 hydrophobic pocket** (uploaded as mp4 video file)

## Expanded View Code

A detailed description of the R code used for microarray analysis is uploaded separately.

## Expanded View Tables

**Expanded View table S1:** *In vitro* and *in vivo* studies using crude chitin preparations. Abbreviations: limulus amebocyte lysate assay (LAL), thin layer chromatography (TLC), endotoxin units (EU), intraperitoneal (i.p.), intranasal (i.n.), murine (M), human (H).

| Publication or study | Nature of chitin used | System | Application | Information regarding purity | Degree of acetylation |
| --- | --- | --- | --- | --- | --- |
| Reese Nature 2007 | Colloidal chitin (NEB), Largest aggregates removed by sedimentation | M | *In vitro*, i.n., i.p. | No information given | Unknown |
| Kigoso J Leukoc Biol 2011 | Colloidal chitin (NEB) >40 µm, Cytometrically sized chitin 1-10 µm | M | *In vitro*, i.p. | Endotoxin removal using Zetapore membrane confirmed by LAL assay | Unknown |
| Wagner Eur J Immunol 2010 | Colloidal chitin from crab shell (Sigma) Largest aggregates removed by sedimentation | M | *In vitro*, i.n. | E-toxate test to exclude LPS | Unknown |
| Wagener PLoS Path 2014 | Crude extraction from *C. albicans* NGY152, Cytometrically sized chitin 1-10 µm | M and H cells | *In vitro*, i.p. | 98% purity (HPLC) | 55-100% (97%) |
| Rizzetto J Biol Chem 2016 | Chitin from *C. albicans* UC820 | M and H cells | *In vitro*, i.p. | 98% purity (HPLC) | Unknown |
| Khosravi PLoS One 2016 | Chitin from *M. rouxii*, Size ~40 µm | H cells | *In vitro* | Endotoxin ≤ 0.1 EU/ml (LAL) | Unknown |
| Becker MBio 2016 | Chitin from *A. fumigatus*, 70% of particles <0.5 µm | H cells | *In vitro* | Removal of β-(1,3)-glucan | 91% |
| Sigsgaard In Forum Allergy Rhinol 2015 | Chitin from *P. borealis* prawn, Milled to 1-20 µm particles | H subjects | I.n. | Endotoxin ≤ 1 EU/ml (LAL) | Unknown |
| Shen Int J Mol Sci 2015 | Chitin and oligo-chitin (no further information given) | M | I.p. | Unknown | Unknown |
| Dubey Immunobiol 2014 | Chitin from crab shell, Filtered through 40 µm strainer, Cytometrically sized chitin 1-10 µm | M | I.p. | Endotoxin ≤ detection (LAL) | Unknown |
| Zakzuk In Arch Allergy Immunol 2014 | Chitin from crab shell (Sigma), High molecular fraction prepared by centrifugation or 40-70 µm fraction | M | I.p., i.n. | Unknown | Unknown |
| Van Dyken Immunity 2014 | Chitin beads (NEB), Filtration 50-70 µm | M | I.p., i.n. | Unknown | Unknown |
| O'Dea Infect Immun 2014 | *A. fumigatus* conidia | M | I.n. | Unknown | Unknown |
| Huang H Immunol 2014 | See Kogiso et al |  |  |  |  |
| Dubey Immunobiology 2014 | Chitin beads (NEB) and *A. fumigatus* preparation 40-µm filtered, cytometrically sized chitin 1-10 µm | M | I.n. | Removal of β-(1,3)-glucan<br>Endotoxin ≤ detection (LAL) | Unknown |
| Alvarez Molecules 2014 | Chitin from *C. albicans* NGY152, *A. fumigatus* 237 and other strains, Crab shell chitin (Sigma), Cytometrically sized around 0.2 µm | H cells | *In vitro* | Unknown | >90% |
| Libreros Int J Cancer 2012 | Chitin 1-10 µm as in publications by Shibata lab | M | *In vitro*, i.p. | Unknown | Unknown |
| Bueter J Biol Chem 2011 | Crude chitosan and chitin (Primex), chitosan and chitin hexamers* | M | *In vitro* | Unknown | Chitosan 76%<br>deacetylated, chitin 93%<br>(TLC) |
| Mora-Montes Infect Immun 2011 | Chitin from *C. albicans* strains | M and H cells | *In vitro* | protein, mannan, and glucan free | 95% |
| Baaten Vaccine 2010 | Chitin particles from shrimps, milled to mean 3.72 µm | M | I.n. | Endotoxin <50 EU/g | Unknown |
| Da Silva J Immunol 2008 | Chitin powder (Sigma) sonicated, filtered and separated into 70-100, 40-70 and <40 µm fractions | M | *In vitro*, i.n. | Endotoxin ≤ detection (LAL) | Unknown |
| Shibata Infect Immunity 1997 | Chitin powder (Sigma) sonicated, Cytometrically sized to 1-10 and 50-100 µm | M | *In vitro*, i.v. | Unknown | Unknown |
| Ozdemir Clin Experimental Allergy 2006 | Chitin microparticles (CMP Therapeutics) from fresh water shrimp, milled to average 4 µm | M | I.n. | Endotoxin <50 EU/g | Unknown |

\* hexamers only used *in vitro* in 1 experiment regarding NLRP3 inflammasome activity.

### Expanded View table S2: Proteomics screen dataset and KEGG pathway mapping

(uploaded as Excel file and submitted to PRIDE online repository)

### Expanded View table S3: Microarray dataset and chitin signature genes

(uploaded as Excel file and submitted to GEO online repository)

**Expanded View table S4:**
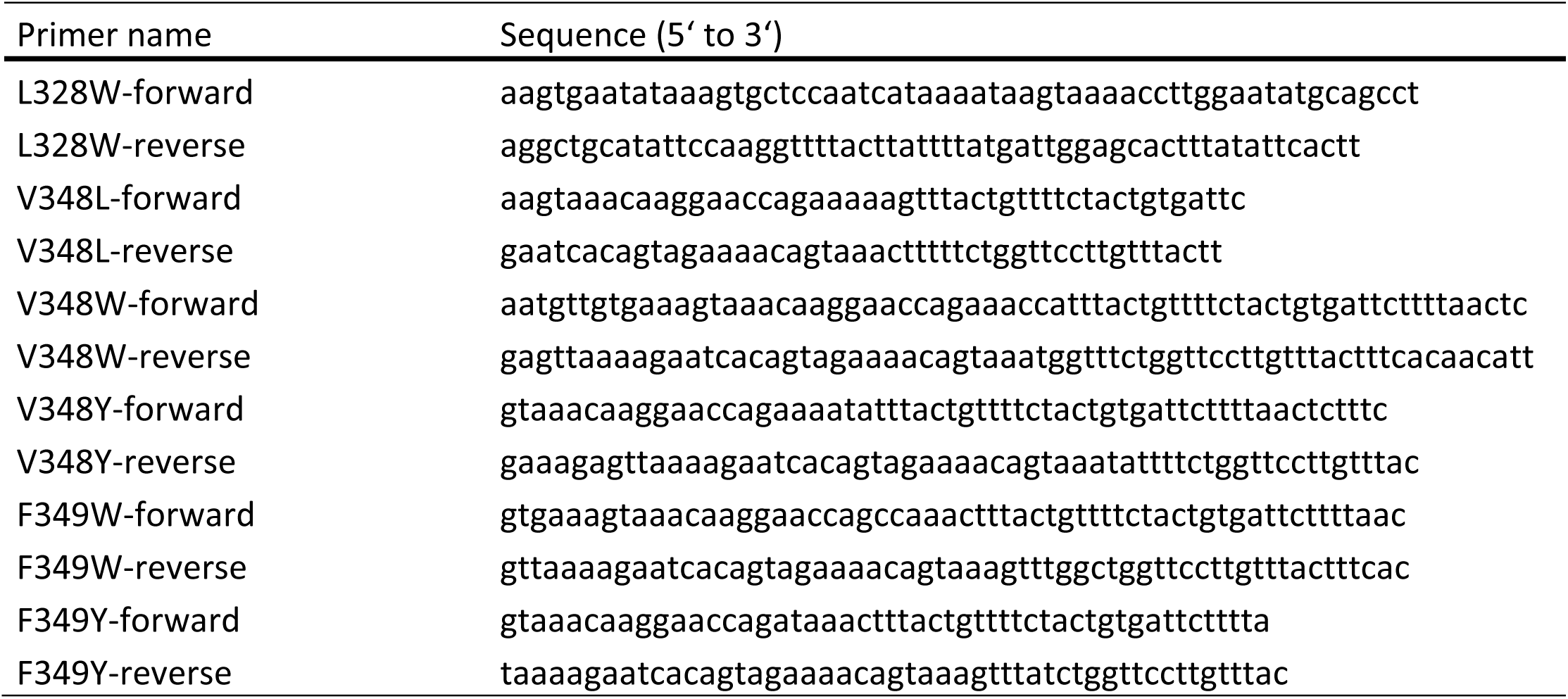
Mutagenesis primers.

**Expanded View table S5:** TaqMan Primers.

| Gene | Assay ID | DNA-Digest |  |
| --- | --- | --- | --- |
|  |  | Yes | No |
| *CAMP* | Hs00189038_m1 |  | X |
| *CCL3* | Hs00234142_m1 |  | X |
| *IFNB1* | Hs01077958_s1 | X |  |
| *IL10* | HS00961622_m1 |  | X |
| *IL12B* | Hs01011518_m1 |  | X |
| *IL1B* | Hs01555410_m1 |  | X |
| *IL6* | Hs00985639_m1 |  | X |
| *IL8* | Hs00174103_m1 |  | X |
| *TBP* | HS00427620_m1 |  | X |

**Expanded View table S6:** siRNAs.

| siRNA | Cat. No. |
| --- | --- |
| siGENOME Non-Targeting Pool#1 | D-001206-13-05 |
| siGENOME H TLR2 (7097) siRNA -SMART pool | M-005120-03-005 |
| ON-TARGETplus MYD88 siRNA - SMARTpool: | L-004769-00-0005 |

## R Code Supplement

### R code to process Whole Blood Microarrays after Stimulation (chitin and others)

This report uses various R packages to walk the reader from the raw data to the paper’s main figures. It is intended to create full transparency and computational reproducibility, and further offers deeper insights into the data than possible in the main figure. We welcome any feedback and invite you to keep it friendly and constructive.

This report was compiled by Felix Frauhammer (whose personal opinion is that all published papers should include the code of their computational analysis) using R function knitr::spin(“file",knit=F) and RStu-dio’s Compile Report button to knit the Rmd into PDF.

date() *# date of compilation*

`## [1] “Mon Mar 5 14:47:18 2018”`

This report goes together with the following paper:

- Whole blood gene expression after stimulation with TLR2 ligands (chitin oligomers, Pam2, Pam3) and the TLR4-ligand LPS
- Corresponding author: Alexander NR Weber
- GEO record GSE103094 (download the GSE103094_non-normalized.txt.gz and unzip it to get started)

`#`

### Information on the 24 microarrays in this dataset

It has the following treatments:

- Unstimulated
- PAM2: Pam2CSK4 (TLR2 ligand)
- PAM3: Pam3CSK4 (TLR2 ligand)
- Chit: chitin oligomers (object of this study)
- LPS: lipopolysaccharide (TLR4 ligand)

`#`

It has the following donors:

- P1: 4 treatments
- P2: 4 treatments
- P3: 5 treatments
- P4: 5 treatments
- P5: 5 treatments

`#`

Install these packages if not already available on your machine:

~~~
**library**(limma)
**library**(ggplot2)
**library**(cowplot)
**library**(tidyverse)
**library**(pheatmap)
**library**(ComplexHeatmap)
*# see end of script for the package versions.*
~~~

## Read in data

### Option1: raw IDAT files

While we recommend using the next paragraph (“Option2: GSE103094_non-normalized.txt”), here are useful hints as to how you would process the raw files.

This data was generated using the **’Illumina Arrays HumanHT-12v4’**, and the Illumina BeadScan software outputs IDAT-files. In order to read these into R, you need to download the ∼ 70 MB Manifest File “HumanHT-12 v4.0 Manifest File (TXT Format)” from the chip’s product files page, next to the raw array files from GEO, and then do this:

~~~
*# idatfiles.all <-dir(pattern="idat”) # in download directory*
*# #idatfiles.all should now be a character vector with IDAT-filepaths*
*# chitin.raw <-limma::read.idat(idatfiles.all, bgxfile="path/to/manifest-file.bgx”)*
*# #should produce the same as ‘chitin.raw’ produced in next paragraph*
~~~

### Option2: GSE103094_non-normalized.txt

We assume you have downloaded and unzipped the supplementary file from our GEO Record. Then you can read it into R like this:

~~~
fileFromGeo <-“/home/felix/chitin_microarrays/data/GSE103094_non-normalized.txt”
*# adapt path to where you downloaded the file*
GEOdata <-**read.csv**(fileFromGeo, header=T, sep="\t”)
~~~

Quick overview over this table:

- The Column “Status” lists the control probes (ERCCs, negative controls, etc.) and then the 47323 probes measuring gene expression.
- each sample has two columns, one with gene expression and one with the DetectionPvalue, i.e. whether the gene in this array is significantly above background signal
- The other three columns are Illumina “Probe_Id", “Array_Address_Id” and, most useful for the biology, the Gene Symbol.

`#`

### Convert to limma class “EListRaw”

~~~
*# extract expression columns:*
geneExprs <-**as.matrix**(GEOdata[,-**c**(1:4)])
geneExprs <-geneExprs[,!**grepl**(“Detection", **colnames**(geneExprs))]
*# Create EListRaw object. Note that the neqc function below uses the information in*
*# column ‘Status’ of the the genes-slot to know which probes are “negative” and which are*
*# “regular” (i.e. measured Genes) for background correction, etc..*
chitin.raw <-**new**(“EListRaw",**list**(E=geneExprs, genes=GEOdata[,1:4], other=**list**(Detection=GEOdata[,**grepl**(“Detection", **colnames**(GEOdata))])))
~~~

## Normalization, filtering and sample Information

We use limma’s robust neqc-function for normalization, which completes the following steps for us:

- corrects background (as measured by negative probes, provided in slot chitin.raw*genes*Status)
- removes control probes (and the now obsolete slot genes$Status)
- log2-transforms data (see further: limma::limmaUsersGuide(), section 17.3)

~~~
chitin <-**neqc**(chitin.raw)
~~~

Next, we filter the measured genes by removing non-expressed probes. **We** only keep genes detected significantly (p<.05) above background in more than 3 arrays:

~~~
expressed <-**rowSums**(chitin$other$Detection < 0.05) > 3
chitin <-chitin[expressed,] *# filters out ∼ 24,000 probes*
~~~

For analysis below, we want to know the Treatment and Patient identiy of each array. It’s part of the colnames, so we can extract it conveniently here:

~~~
meta <-**data.frame**(**t**(**rbind.data.frame**(**strsplit**(**colnames**(chitin), “_”))),
row.names=**colnames**(chitin))
**colnames**(meta) <-**c**(“Treatment","Patient”)
~~~

## Principal Component Analysis

Below we compute principal component analysis on all non-filtered probes (22929 genes) and then again on 6191 genes that vary significantly between treatment groups according to ANOVA (done with limma’s F-test). In the paper’s main figure, we show the latter as the former is influenced more by patient-patient heterogeneity, as we’ll see below.

`#`

### F-test for high-variance genes

This paragraph identifies genes that vary significantly across treatment groups (Chitin, LPS, PAM2, PAM3, Unstimulated). The 6191 identified genes and their expression values (see object anovaexprs at end of paragraph) will go into PCA, see next heading.

~~~
*# design matrix for lmFit*
ct <-**factor**(meta$Treatment)
*# It is good practice to set the control as first level:*
ct <-**relevel**(ct, “Unstimulated”) design <-**model.matrix**(∼0+ct) **colnames**(design) <- **levels**(ct)
*# duplicate correlation for lmFit*
dupcor <- **duplicateCorrelation**(chitin, design, block=meta$Patient) dupcor$consensus.correlation *# can be fed into lmFit function*
## [1] 0.3112271
~~~

We see that differently treated samples from the same patient correlate moderately with each other (∼0.3), as expected from human material. Notifying lmFit of this patient-patient heterogeneity will make the analysis more robust.

~~~
*# fit limma model:*
fit <- **lmFit**(chitin, design, block=meta$Patient,
             correlation = dupcor$consensus.correlation)
contrasts.all <- **makeContrasts**(
        Chitin-LPS,Chitin-PAM2, Chitin-PAM3, Chitin-Unstimulated,
        LPS-PAM2, LPS-PAM3, LPS-Unstimulated,
        PAM2-PAM3,PAM2-Unstimulated, PAM3-Unstimulated,
        levels=design
)
fit2.all <- **contrasts.fit**(fit, contrasts.all)
fit2.all <- **eBayes**(fit2.all, trend=T)
~~~

The two lines above compute an F-test, which is equivalent to a one-way ANOVA for each gene except that the residual mean squares have been moderated between genes. (see limma::limmaUsersGuide()) After multiple testing correction with Benjamini-Hochberg, 6191 genes are found to vary across different groups with FDR<.05:

~~~
anova <- **topTableF**(fit2.all, p.value=0.05, number=50000, adjust.method = ^1^BH^1^)
**dim**(anova)
## [1] 6191 17
~~~

These genes contain the information on what separates different treatments independent of the patient of origin. To analyze them in PCA, we extract the expression values of these genes:

~~~
anovaexprs <- chitin$E
**rownames**(anovaexprs) <- chitin$genes$Probe_Id
anovaexprs <- anovaexprs[**rownames**(anovaexprs) %in% anova$Probe_Id,]
~~~

## PCA on all genes separates Patients

Human material often shows high patient-patient variability. In line with this, the largest variation captured by PCA on all genes is driven by patient-patient differences, as we see in the following.

~~~
pca.allGenes <- **prcomp**(**t**(chitin$E))
scores.allGenes <- **cbind**(**as.data.frame**(pca.allGenes$x), meta)
screeplot(pca.allGenes, type = “lines”)
~~~

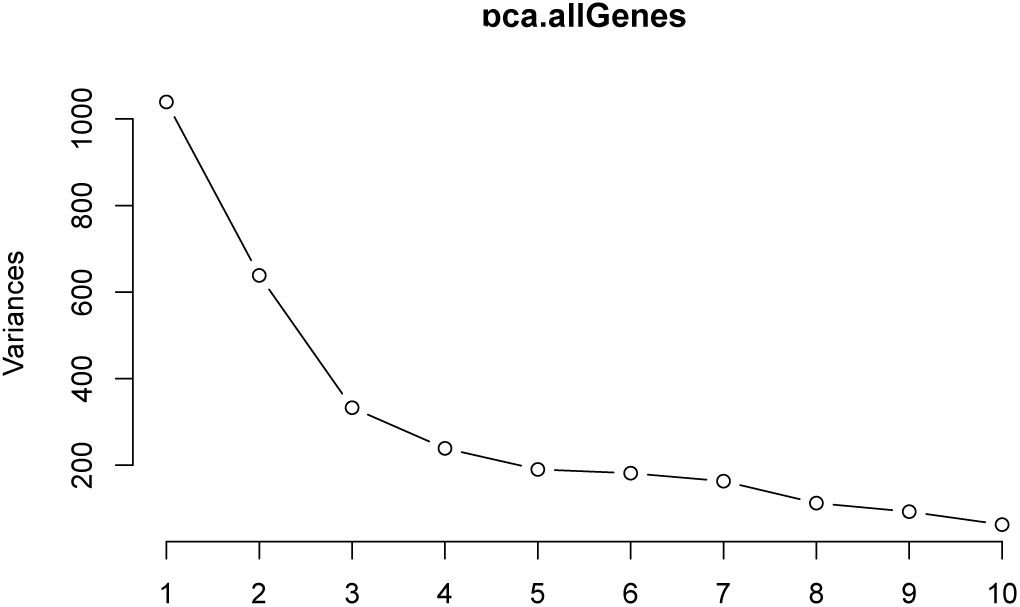

~~~
**ggplot**(scores.allGenes, **aes**(x=PC1, y=PC2, col=Treatment, shape=Patient))+
  geom_point(size=3)+ggtitle(“allGenes”)
~~~

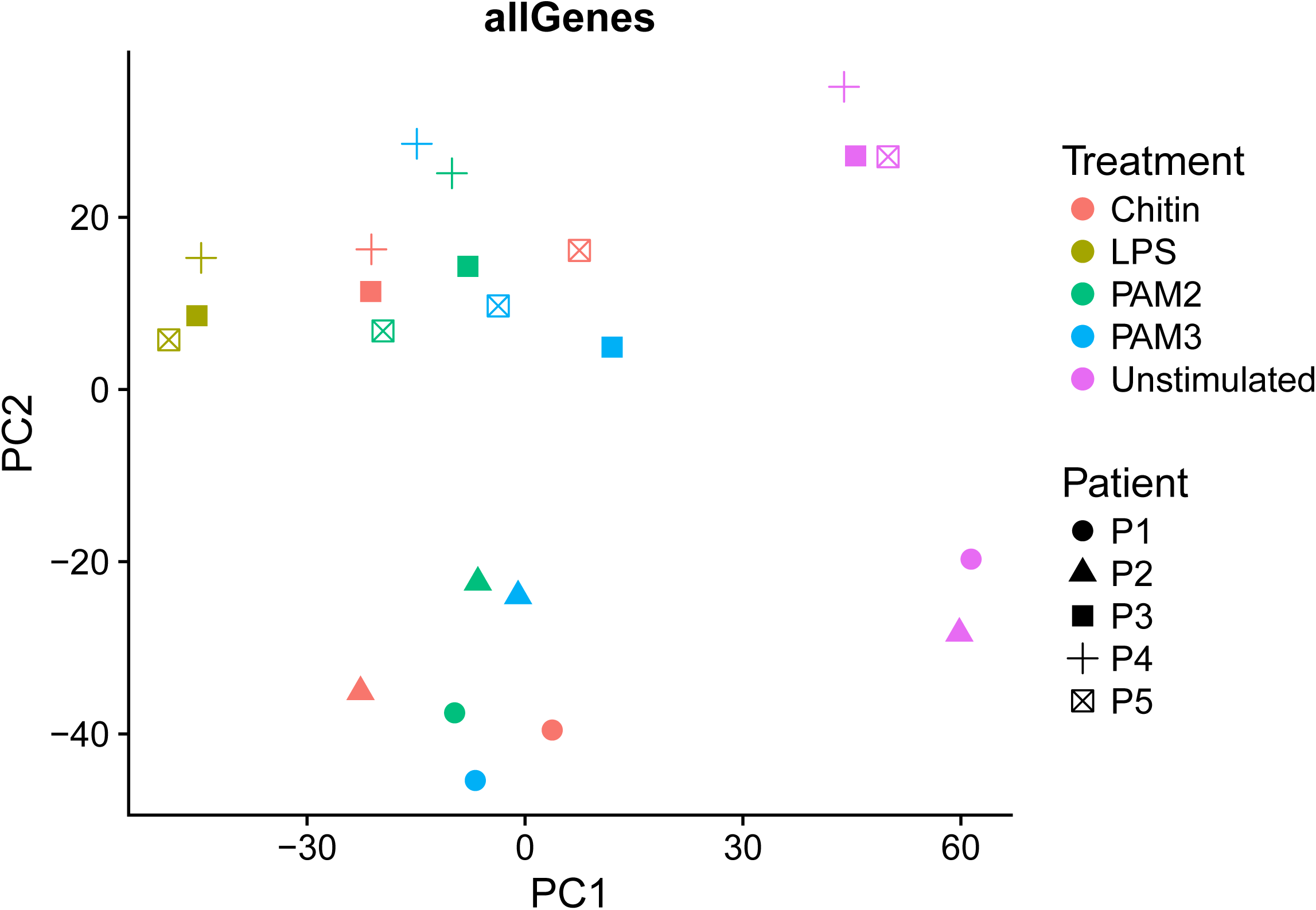

We see that when computed on all genes, PC1 indicates a strong difference between Unstimulated, LPS and the others, while PC2 separates Patients P1 and P2 from other donors. To better resolve the differences of Chitin, PAM2 and PAM3, we compute PCA on genes that vary between groups in the next paragraph, hoping to reduce how much PCA is influenced by donor heterogeneity.

`#`

### PCA on 6191 ANOVA genes

pca.anova<- **prcomp**(**t**(anovaexprs))

scores.anova <- **cbind**(**as.data.frame**(pca.anova$x), meta)

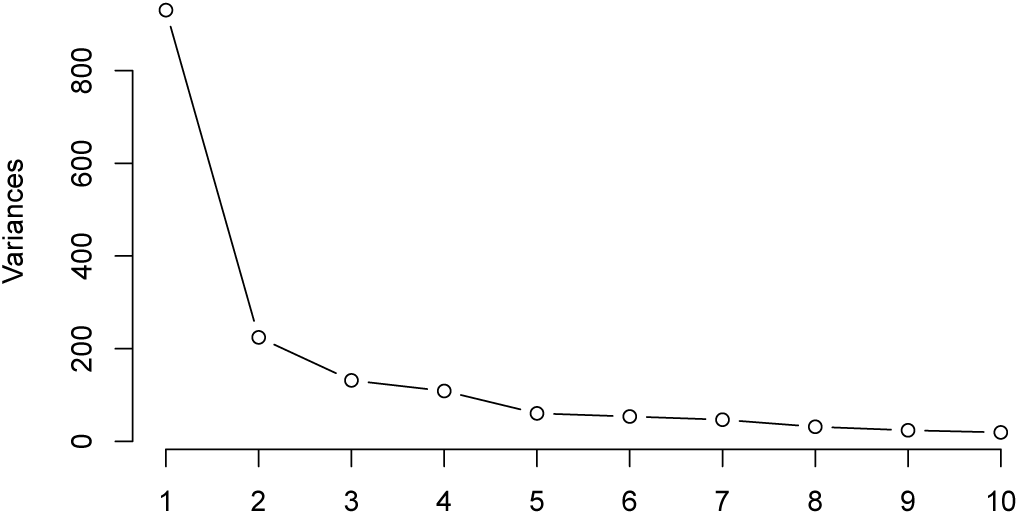

**screeplot**(pca.anova, type = “lines”)

**ggplot**(scores.anova, **aes**(x=PC1, y=PC2, col=Treatment, shape=Patient))+

**geom_point**(size=3)+**ggtitle**(“ANOVA”)

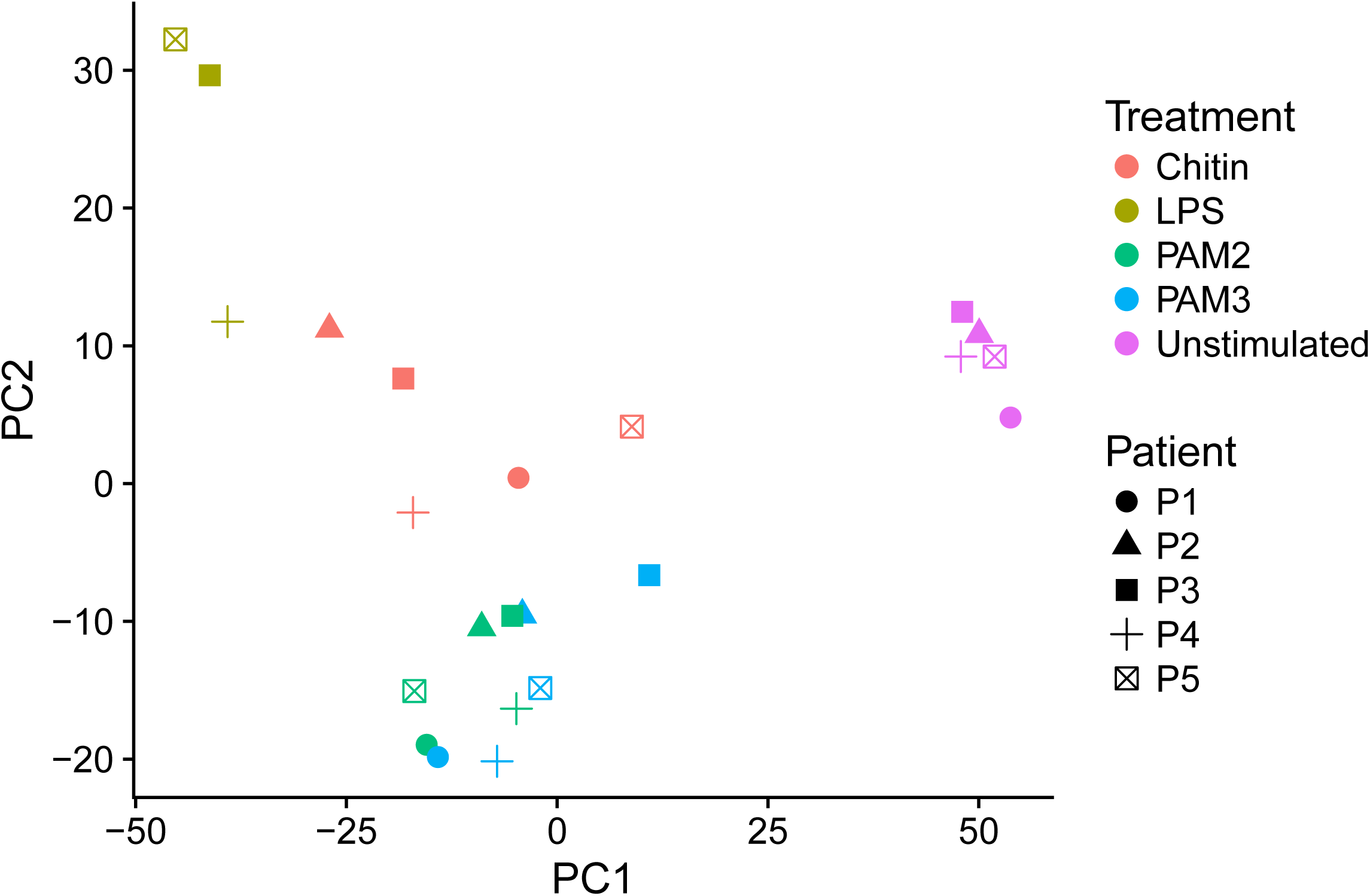

As hoped, the samples do not cluster by patient anymore. As above, PC1 separates Unstimulated and LPS from the others. Interestingly, PAM2 and PAM3 seem very similar to each other, while all chitin samples appear clearly distinct from PAM and LPS along PC2. We note, however, that above scree-plot shows PC2 to play a rather minor role, as most variance in the data is captured by PC1. This already indicates that chitin, PAM2 and PAM3 show rather minor differences relative to the effects induced by LPS. For biological interpretation of this PCA, we will look into the genes contributing to PCs 1 and 2 in the next subparagraph, while the next big heading will find and display Differentially Expressed Genes between treatment groups.

`#`

### Obtain and export PCA loadings

To place the PCA figure in above’s paragraph into biological context, we extract the information on which genes contribute the most to PC1 and PC2.

~~~
**par**(mfrow=**c**(1,2))
**plot**(pca.anova$rotation[,1][**order**(**abs**(pca.anova$rotation[,1]), decreasing=T)],
     main="PC1 loads", ylab="Gene1s contribution (1Load1) to PC1", xlab="Genes”)
**plot**(pca.anova$rotation[,2][**order**(**abs**(pca.anova$rotation[,2]), decreasing=T)],
     main="PC2 loads", ylab="Gene1s contribution (1Load1) to PC2", xlab="Genes”)
~~~

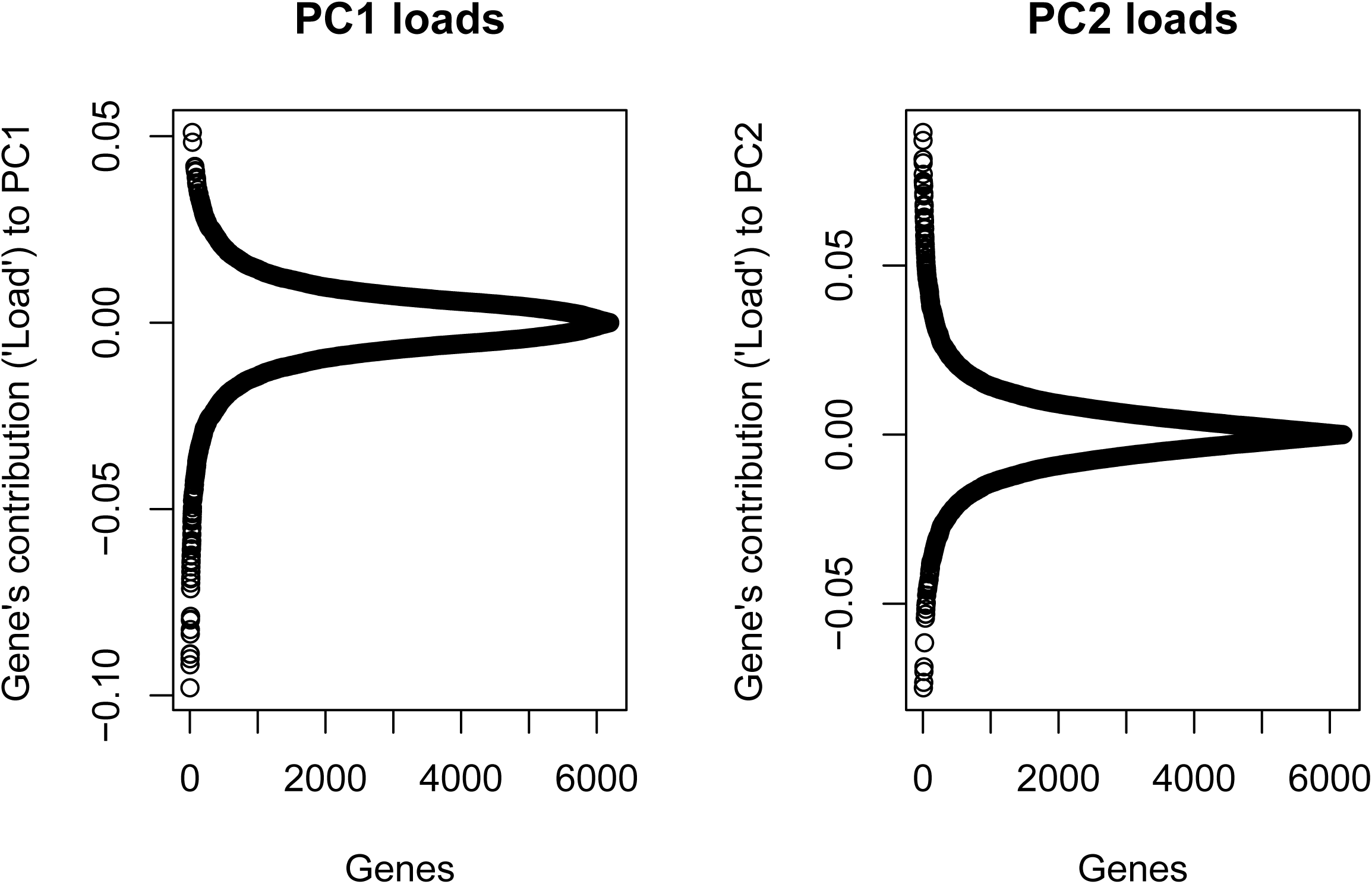

**par**(mfrow=**c**(1,1))

Notes on above plot: Genes with large absolute loads contribute most to PCs. For example, the genes with loads around 0.05 make a sample’s PC1 score positive (e.g. all Unstimulated samples in the “ANOVA"-PCA plot above), whereas high expression of the negative-load genes makes a sample show negative PC1 values (e.g. LPS samples in above plot).

From the plots above we can estimate roughly by eye that perhaps the 100 or 300 genes with the highest load are interesting for biological interpretation, as it is mainly their cumulative load separating the different treatments from each other in PCA plot “ANOVA". Still, we export all genes that went into the PCA, for completeness:

~~~
topPC1 <- pca.anova$rotation[,1, drop=F]
topPC1 <- topPC1[**order**(**abs**(topPC1), decreasing=T),,drop=F]
topPC2 <- pca.anova$rotation[,2, drop=F]
topPC2 <- topPC2[**order**(**abs**(topPC2), decreasing=T),,drop=F]
*# bring it all into same dataframe and add gene Symbol annotations:*
PCtopDF <- **data.frame**(PC1_load=topPC1[,1], PC1_ProbeID=**rownames**(topPC1),
           PC1_Symbol=GEOdata$Symbol[**match**(**rownames**(topPC1), GEOdata$Probe_Id)],
           PC2_load=topPC2[,1], PC2_ProbeID=**rownames**(topPC2),
           PC2_Symbol=GEOdata$Symbol[**match**(**rownames**(topPC2), GEOdata$Probe_Id)],
           stringsAsFactors = F
           )
*# for duplicated symbols (whenever the array measures genes with >1 probes), we*
*# only keep the first probe to avoid confusion.*
PCtopDF <- PCtopDF[!**duplicated**(PCtopDF$PC1_Symbol),]
PCtopDF <- PCtopDF[!**duplicated**(PCtopDF$PC2_Symbol),]
**head**(PCtopDF)
##                PC1_load PC1_ProbeID PC1_Symbol PC2_load PC2_ProbeID
##ILMN_1699651 -0.09794445 ILMN_1699651 IL6 0.08933929 ILMN_1707695
## ILMN_1671509 -0.09177125 ILMN_1671509 CCL3 0.08698045 ILMN_2054019
## ILMN_1747355 -0.09014691 ILMN_1747355 CCL3L1 0.08145545 ILMN_1701789
## ILMN_1805410 -0.08884788 ILMN_1805410 C15orf48 0.07698123 ILMN_1674811
## ILMN_1838319 -0.08351726 ILMN_1838319 LOC730249 0.07488550 ILMN_2067890
## ILMN_1720048 -0.08231378 ILMN_1720048 CCL2 -0.07485910 ILMN_1686116
##              PC2_Symbol
## ILMN_1699651      IFIT1
## ILMN_1671509      ISG15
## ILMN_1747355      IFIT3
## ILMN_1805410       OASL
## ILMN_1838319     CXCL11
## ILMN_1720048      THBS1
~~~

The first 3 columns rank genes according to their influence (load) on PC1, the last 3 for PC2. We now write out the complete table and provide it as supplementary table attached to this paper. Note that it includes all genes that vary across conditions (according to ANOVA), and the first few hundred genes in each column contribute the most to PCs 1 and 2.

~~~
**write_csv**(x = PCtopDF, path = **paste0**(“/home/felix/chitin_microarrays/PC1-2_topGenes",
                                     **format**(**Sys.time**(), “_%Y%m%d-%H.%M”),".csv”))
~~~

## Heatmap for the main figure

The heatmap in the main figure is generated at the end of this section and meant to illustrate and emphasise the existing differences between Chitin and other treatments. It is, however, important to note at this point that in fact most genes up/down-regulated by one treatment are up/down-regulated in most other treatments as well. This is expected, as all treatments activate large immune signaling cascades. To make this fact clear to the reader, we now quickly compare the log-foldchanges of different treatments with each other, before moving on to the main heatmap.

~~~
*# get logFCs for each sample (“relative expression", relExprs)*
relExprs <-NULL
for(patient in levels(meta$Patient)){ relExprs <-cbind(relExprs, apply(chitin$E[,
grepl(patient, colnames(chitin$E))],2, function(column) column-chitin$E[,paste0(“Unstimulated_",patient)]))
}
relExprs <- relExprs[,!grepl(“Unstimulated", colnames(relExprs))]
*# average logFCs per treatment:*
meanLogFC <- NULL
Ts <-c(“Chitin","LPS", “PAM2","PAM3”)
for(treatment in Ts){
meanLogFC <- cbind(meanLogFC,
rowMeans(relExprs[,grepl(treatment, colnames(relExprs))]))
}
colnames(meanLogFC) <- Ts
gather(as.data.frame(meanLogFC),key="Treatment", value="MeanLogFC", LPS:PAM3) %>%
ggplot(aes(x=Chitin, y=MeanLogFC))+geom_point()+facet_wrap(∼Treatment)+
xlab(“Mean logFC of Chitin samples”)+ geom_smooth(method =lm)
~~~

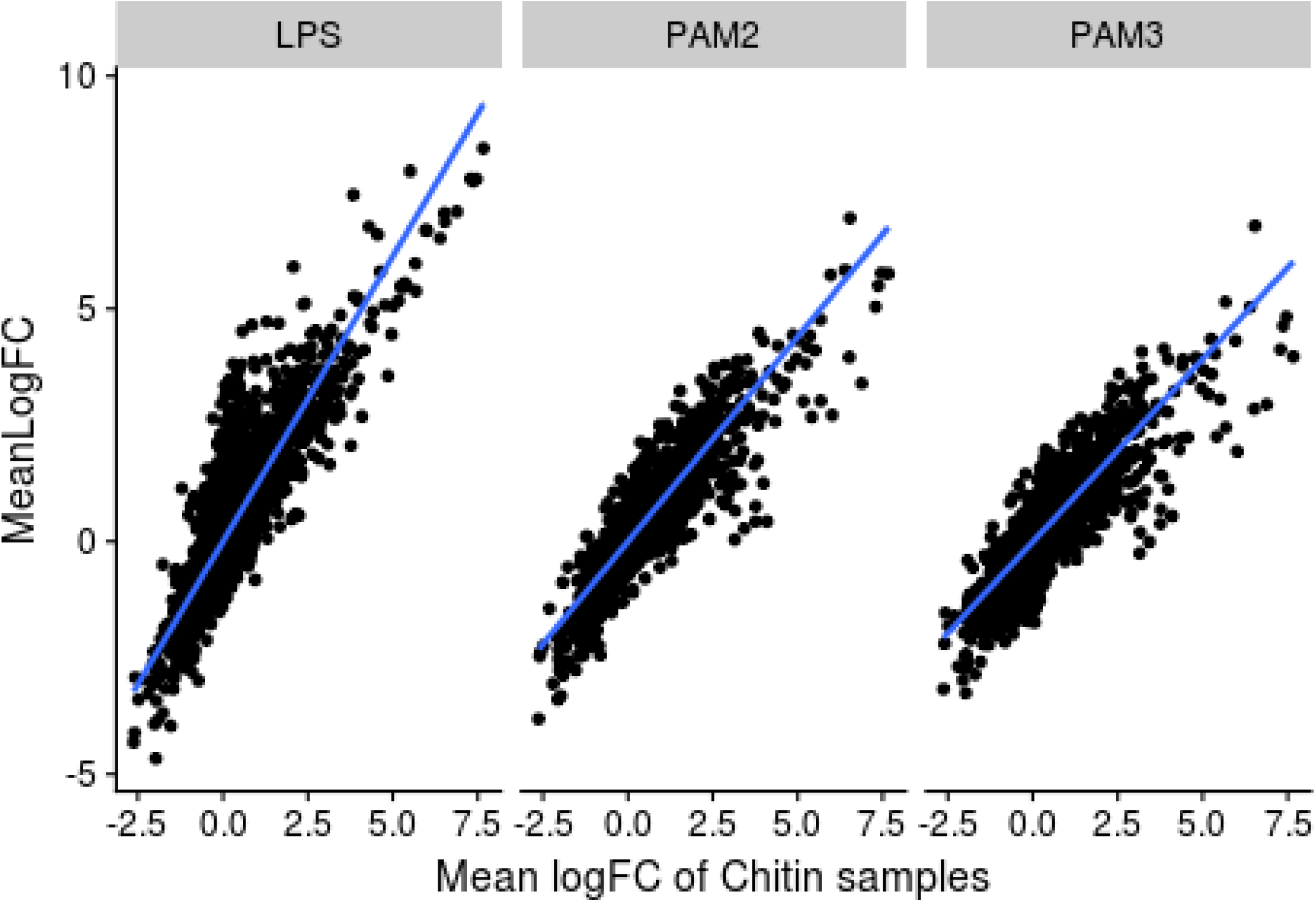

As we see, whenever chitin-treatment induces a high log-foldchange for a gene, the other treatments do also lead to upregulation. One thing to note is that LPS has particularly many upregulations that do not change for chitin samples (left panel, note the many dots at x=0 lying far above the blue line), and LPS also has the steepest line, indicating all genes are upregulated more than for other treatments. Again, this is not unexpected, as we activate strong immune responses with all treatments. Still, we wanted to make this point clear so that the heatmap in the main paper figure can be read in the right context.

`#`

## Find chitin-specific genes

Now that we are aware how all treatments induce similar big transcriptional changes, it is interesting to work out differences between them. For this, we will compute differentially expressed genes between Chitin and the other treatments, and display them in a heatmap for the main paper.

~~~
*# chitin-specific genes (higher/lower than all other treatments)*
contrasts.cs <- **makeContrasts**(
Chitin-PAM2,Chitin-PAM3, Chitin-LPS,
levels=design)
~~~

Firstly, we are interested in genes where chitin is expressed highest. These are not too many, as we just saw that LPS is the most potent inducer of gene expression, so we choose a rather loose cutoff of foldchange=1.1 to be able to look at a few:

~~~
lfc1.1 <- **decideTests**(**treat**(**contrasts.fit**(fit, contrasts.cs), lfc = **log2**(1.1)))
chit_gt_all <- chitin[**rowSums**(lfc1.1)==3,]
**dim**(chit_gt_all)
## [1] 11 23
~~~

11 genes are 10 % or more higher in chitin than all three other treatments.

`#`

Other than being strictly highest, we next ask for genes where Chitin is the most different from the other Treatments. For this we identify genes that show misregulation compared to both PAMs (foldchange 1.2 or higher), irrespective of whether it is up- or downregulated. For a better overview in the final heatmap, we choose to split the resulting genes into two groups: one where chitin and LPS show no sign. difference (chit_LPSnoChange) and one where they do (chit_LPS_DE).

~~~
lfc1.2<- **decideTests**(**treat**(**contrasts.fit**(fit, contrasts.cs), lfc = **log2**(1.2)))
*# identify genes significantly up- OR downregulated between chitin and PAM2 OR PAM3.*
chit_LPSnoChange <-chitin[ (lfc1.2[,1]!=0 & lfc1.2[,2]!=0) & lfc1.2[,3]==0, ]
chit_LPS_DE <- chitin[(lfc1.2[,1]!=0 & lfc1.2[,2]!=0) & lfc1.2[,3]!=0, ]
*# to avoid confusion, if multiple array probes measured the same gene we only keep one:*
chit_LPSnoChange <- chit_LPSnoChange[!**duplicated**(chit_LPSnoChange$genes$Symbol),]
chit_LPS_DE <- chit_LPS_DE[!**duplicated**(chit_LPS_DE$genes$Symbol),]
*# we also want to display each gene only once in the heatmap:*
chit_LPS_DE <- chit_LPS_DE[! chit_LPS_DE$genes$Symbol %in% chit_gt_all$genes$Symbol,] chit_LPSnoChange <- chit_LPSnoChange[! chit_LPSnoChange$genes$Symbol %in%
chit_gt_all$genes$Symbol,]
~~~

### Create Heatmap from main figure

In the above paragraph we have selected interesting genes to look at, and now display them in a heatmap.

~~~
*# We first sort the columns of the expression matrix by treatments:*
heatmap.E <- **data.frame**(Symbol=chitin$genes$Symbol,
   chitin$E[,**grepl**(“Unstimulated", **colnames**(chitin$E))],
   chitin$E[,**grepl**(“PAM3",**colnames**(chitin$E))],
   chitin$E[,**grepl**(“PAM2", **colnames**(chitin$E))],
   chitin$E[,**grepl**(“Chitin",**colnames**(chitin$E))],
   chitin$E[,**grepl**(“LPS", **colnames**(chitin$E))])
  **rownames**(heatmap.E) <- chitin$genes$Probe_Id
*# we now extract the genes of interest, standardize them with ‘scale’ (to get # z-scores) and assign the corresponding gene Symbols:*
genesOfInterest<-**c**(**as.character**(chit_gt_all$genes$Probe_Id),
                   **as.character**(chit_LPS_DE$genes$Probe_Id),
                   **as.character**(chit_LPSnoChange$genes$Probe_Id))
hm <- heatmap.E[**match**(genesOfInterest, **rownames**(heatmap.E)), ]
**rownames**(hm) <- hm$Symbol; hm$Symbol <- **c**()
hm <- **data.frame**(**t**(**scale**(**t**(hm))))
~~~

We create the heatmap as in the main figure using the ComplexHeatmap package. Note that we deactivate clustering columns on purpose (while clustering is a great approach for unsupervised analysis, using it here would defeat its purpose: the heatmap should visualize genes selected by their differential expression, which is a supervised approach).

~~~
**Heatmap**(hm, cluster_columns = F, cluster_rows = T,
        show_row_dend=F,
                  split=**c**(
                      **rep**(“>all\n logFC>1.1",**dim**(chit_gt_all)[1]),
                      **rep**(“DE: both PAMs & LPS\n logFC>1.2",**dim**(chit_LPS_DE)[1]),
                      **rep**(“DE: both PAMs\n logFC<1.2",**dim**(chit_LPSnoChange)[1])),
       name=**paste**(“z-scores","(Expression)", sep = “\n”), row_names_gp = **gpar**(fontsize=11)
       )
~~~

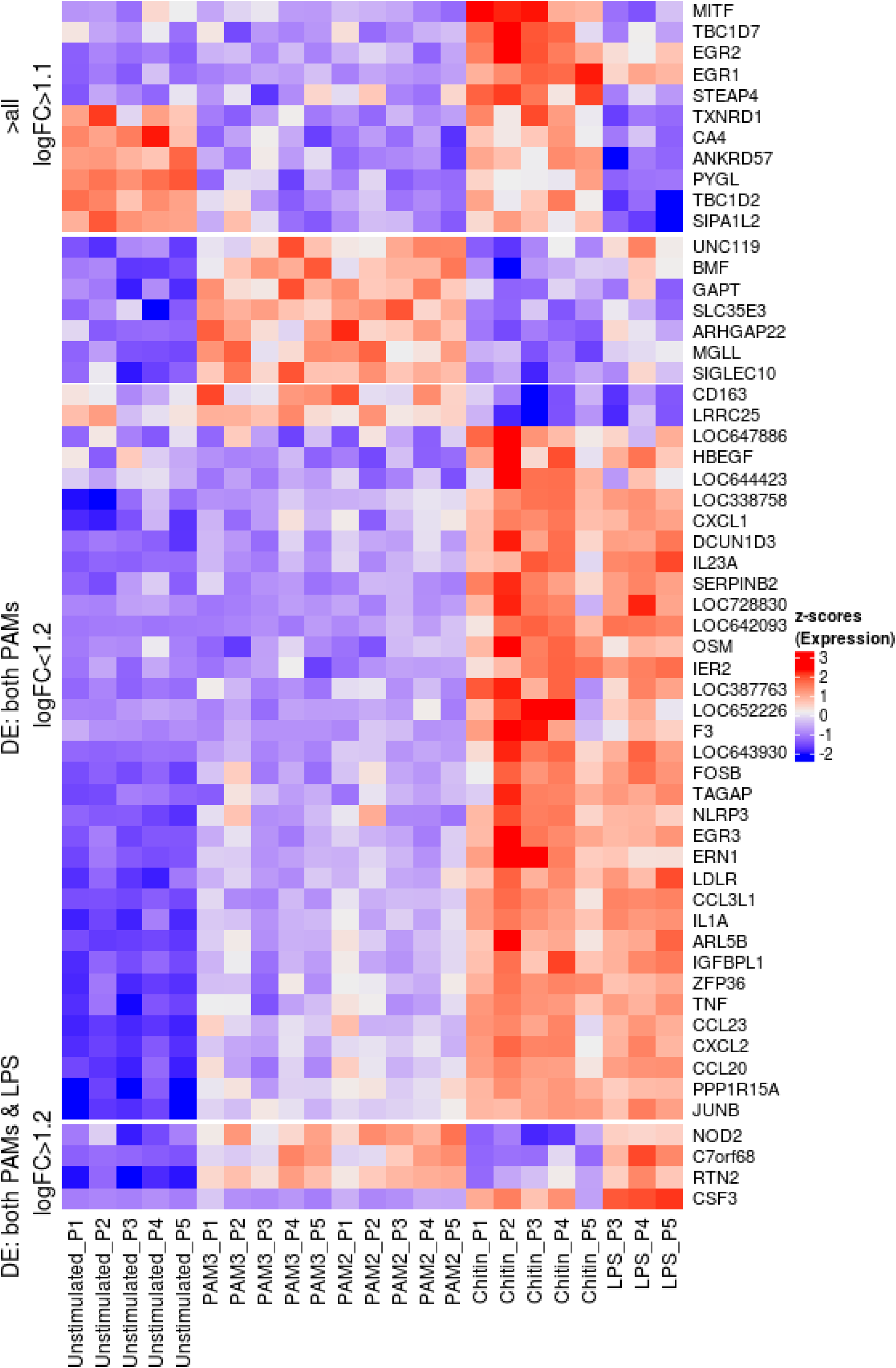

## Supplementary figure: PC top genes

We have used PCA to illustrate treatment differences. Here, we place the PCs into biological context by showing the expression of the 100 genes contributing most to PC1 and PC2.

~~~
hm.pc1 <- heatmap.E[**match**(PCtopDF$PC1_ProbeID[1:100], **rownames**(heatmap.E)), ]
**rownames**(hm.pc1) <- hm.pc1$Symbol; hm.pc1$Symbol <- **c**()
hm.pc1 <- **data.frame**(**t**(**scale**(**t**(hm.pc1))))
**Heatmap**(hm.pc1, cluster_columns = F, cluster_rows=T, name="PC1 Top100", row_names_gp = **gpar**(fontsize=8))
~~~

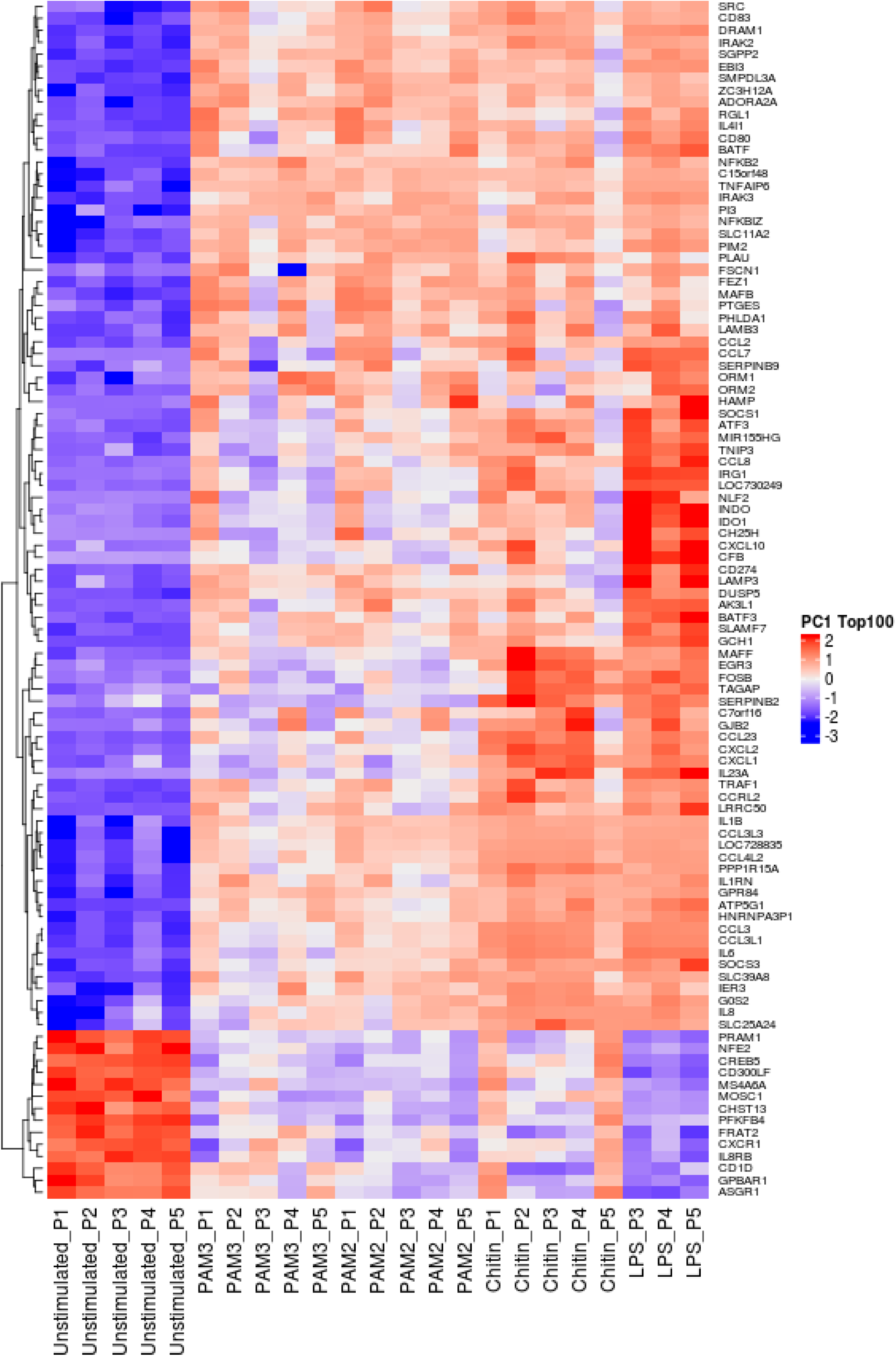

In above heatmap, we observe again what we already saw in the logFC-plot above: most genes show the same qualitative reaction (up or down) for all treatments, but differ by how much (e.g. LPS up/downregulates the most).

~~~
hm.pc2 <- heatmap.E[**match**(PCtopDF$PC2_ProbeID[1:100], **rownames**(heatmap.E)), ]
**rownames**(hm.pc2) <- hm.pc2$Symbol; hm.pc2$Symbol <- **c**()
hm.pc2 <- **data.frame**(**t**(**scale**(**t**(hm.pc2))))
**Heatmap**(hm.pc2, cluster_columns = F, cluster_rows=T, name="PC2 Top100", row_names_gp = **gpar**(fontsize=8))
~~~

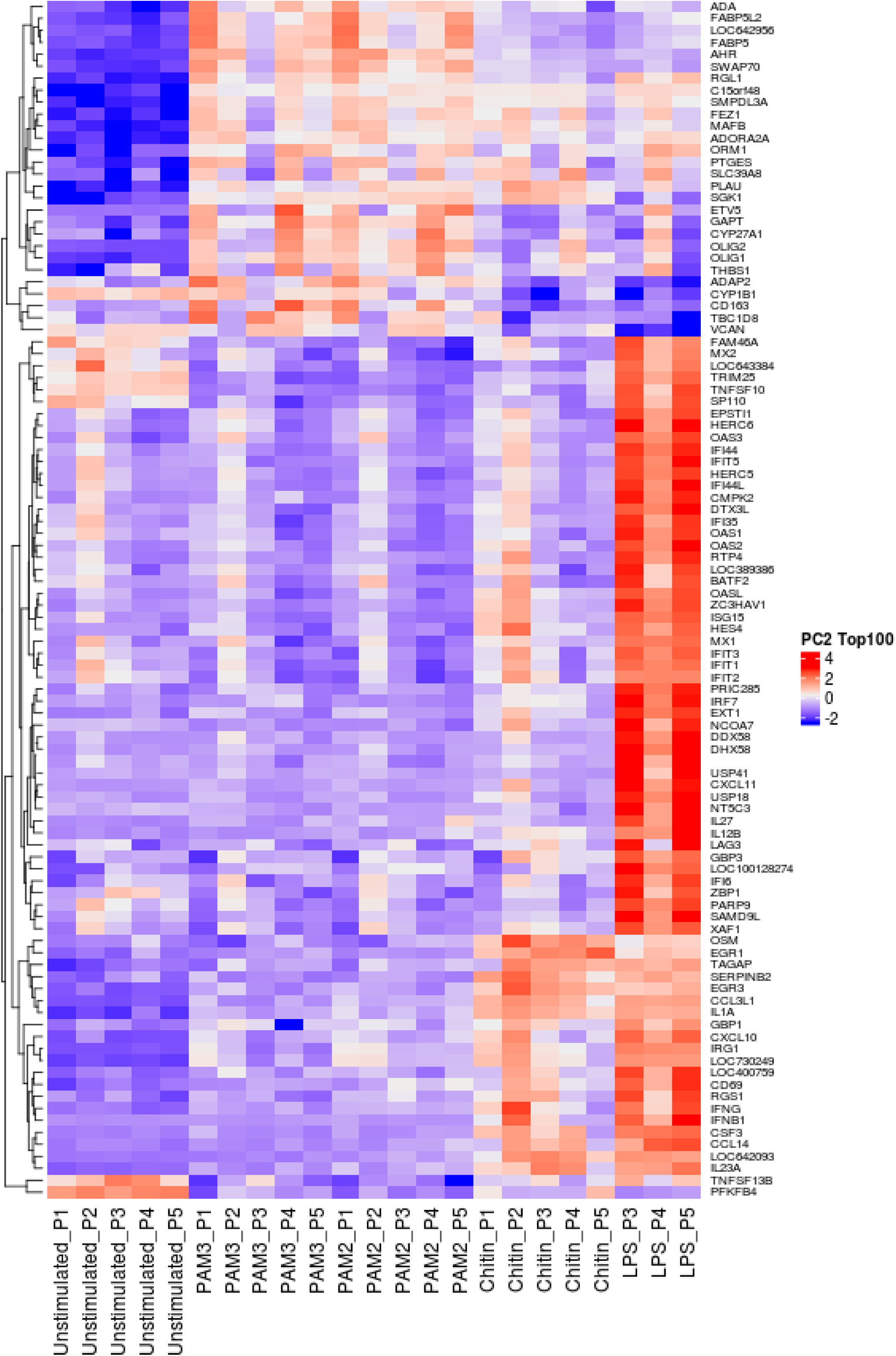

For PC2, we see that LPS has many additional genes upregulated which are almost unchanged in the other treatments. Also, Chitin sometimes behaves more like LPS, and sometimes more like PAMs. We also remark that donor P2 is behaving somewhat as an outlier.

`#`

## End of Script

~~~
**sessionInfo**()
## R version 3.4.2 (2017-09-28)
## Platform: x86_64-pc-linux-gnu (64-bit)
## Running under: Ubuntu 14.04.5 LTS
##
## Matrix products: default
## BLAS: /usr/lib/libblas/libblas.so.3.0
## LAPACK: /usr/lib/lapack/liblapack.so.3.0 ##
## locale:
## [1] LC_CTYPE=en_US.UTF-8 LC_NUMERIC=C
## [3] LC_TIME=de_DE.UTF-8 LC_COLLATE=en_US.UTF-8
## [5] LC_MONETARY=de_DE.UTF-8 LC_MESSAGES=en_US.UTF-8
## [7] LC_PAPER=de_DE.UTF-8 LC_NAME=C
## [9] LC_ADDRESS=C LC_TELEPHONE=C
## [11] LC_MEASUREMENT=de_DE.UTF-8 LC_IDENTIFICATION=C ##
## attached base packages:
## [1] grid stats graphics grDevices utils datasets methods
## [8] base
## other attached packages:
## [1] ComplexHeatmap_1.17.1 pheatmap_1.0.8 forcats_0.2.0
## [4] stringr_1.3.0 dplyr_0.7.4 purrr_0.2.4
## [7] readr_1.1.1 tidyr_0.8.0 tibble_1.4.2
## [10] tidyverse_1.2.1 cowplot_0.9.2 ggplot2_2.2.1
## [13] limma_3.34.8
## loaded via a namespace (and not attached):
~~~

## References

Albert M, Furst U (2017) Quantitative Detection of Oxidative Burst upon Activation of Plant Receptor Kinases. Methods Mol Biol 1621: 69–76

Alvarez FJ (2014) The effect of chitin size, shape, source and purification method on immune recognition. Molecules 19: 4433–4451

Boersema PJ, Raijmakers R, Lemeer S, Mohammed S, Heck AJ (2009) Multiplex peptide stable isotope dimethyl labeling for quantitative proteomics. Nat Protoc 4: 484–494

Brock AK, Willmann R, Kolb D, Grefen L, Lajunen HM, Bethke G, Lee J, Nurnberger T, Gust AA (2010) The Arabidopsis mitogen-activated protein kinase phosphatase PP2C5 affects seed germination, stomatal aperture, and abscisic acid-inducible gene expression. Plant Physiol 153: 1098–1111

Brown GD, Denning DW, Gow NA, Levitz SM, Netea MG, White TC (2012) Hidden killers: human fungal infections. Sci Transl Med 4: 165rv113

Bueter CL, Lee CK, Rathinam VA, Healy GJ, Taron CH, Specht CA, Levitz SM (2011) Chitosan but not chitin activates the inflammasome by a mechanism dependent upon phagocytosis. J Biol Chem 286: 35447–35455

Bueter CL, Specht CA, Levitz SM (2013) Innate sensing of chitin and chitosan. PLoS Pathog 9: e1003080

Cho JS, Guo Y, Ramos RI, Hebroni F, Plaisier SB, Xuan C, Granick JL, Matsushima H, Takashima A, Iwakura Y et al (2012) Neutrophil-derived IL-1beta is sufficient for abscess formation in immunity against Staphylococcus aureus in mice. PLoS Pathog 8: e1003047

Choi JP, Lee SM, Choi HI, Kim MH, Jeon SG, Jang MH, Jee YK, Yang S, Cho YJ, Kim YK (2016) House Dust Mite-Derived Chitin Enhances Th2 Cell Response to Inhaled Allergens, Mainly via a TNF-alpha-Dependent Pathway. Allergy Asthma Immunol Res 8: 362–374

Cox J, Mann M (2008) MaxQuant enables high peptide identification rates, individualized p.p.b.-range mass accuracies and proteome-wide protein quantification. Nat Biotechnol 26: 1367–1372

Cox J, Neuhauser N, Michalski A, Scheltema RA, Olsen JV, Mann M (2011) Andromeda: a peptide search engine integrated into the MaxQuant environment. J Proteome Res 10: 1794–1805

Cunha C, Romani L, Carvalho A (2010) Cracking the Toll-like receptor code in fungal infections. Expert Rev Anti Infect Ther 8: 1121–1137

Da Silva CA, Chalouni C, Williams A, Hartl D, Lee CG, Elias JA (2009) Chitin is a size-dependent regulator of macrophage TNF and IL-10 production. J Immunol 182: 3573–3582

Da Silva CA, Hartl D, Liu W, Lee CG, Elias JA (2008) TLR-2 and IL-17A in chitin-induced macrophage activation and acute inflammation. J Immunol 181: 4279–4286

Di Carlo FJ, Fiore JV (1958) On the composition of zymosan. Science 127: 756–757

Fischer M, Spies-Weisshart B, Schrenk K, Gruhn B, Wittig S, Glaser A, Hochhaus A, Scholl S, Schnetzke U (2016) Polymorphisms of Dectin-1 and TLR2 Predispose to Invasive Fungal Disease in Patients with Acute Myeloid Leukemia. PLoS One 11: e0150632

Gantner BN, Simmons RM, Canavera SJ, Akira S, Underhill DM (2003) Collaborative induction of inflammatory responses by dectin-1 and Toll-like receptor 2. J Exp Med 197: 1107–1117

Goodridge HS, Underhill DM (2008) Fungal Recognition by TLR2 and Dectin-1. Handb Exp Pharmacol: 87–109

Hayafune M, Berisio R, Marchetti R, Silipo A, Kayama M, Desaki Y, Arima S, Squeglia F, Ruggiero A, Tokuyasu K et al (2014) Chitin-induced activation of immune signaling by the rice receptor CEBiP relies on a unique sandwich-type dimerization. Proc Natl Acad Sci U S A 111: E404–413

Hornak V, Abel R, Okur A, Strockbine B, Roitberg A, Simmerling C (2006) Comparison of multiple Amber force fields and development of improved protein backbone parameters. Proteins 65: 712– 725

Hoving JC, Wilson GJ, Brown GD (2014) Signalling C-type lectin receptors, microbial recognition and immunity. Cellular microbiology 16: 185–194

Imai T, Watanabe T, Yui T, Sugiyama J (2002) Directional degradation of beta-chitin by chitinase A1 revealed by a novel reducing end labelling technique. FEBS letters 510: 201–205

Jerabek-Willemsen M, Wienken CJ, Braun D, Baaske P, Duhr S (2011) Molecular interaction studies using microscale thermophoresis. Assay Drug Dev Technol 9: 342–353

Jimenez-Dalmaroni MJ, Radcliffe CM, Harvey DJ, Wormald MR, Verdino P, Ainge GD, Larsen DS, Painter GF, Ulevitch R, Beutler B et al (2015) Soluble human TLR2 ectodomain binds diacylglycerol from microbial lipopeptides and glycolipids. Innate Immun 21: 175–193

Jin MS, Kim SE, Heo JY, Lee ME, Kim HM, Paik SG, Lee H, Lee JO (2007) Crystal structure of the TLR1-TLR2 heterodimer induced by binding of a tri-acylated lipopeptide. Cell 130: 1071–1082

Jones JD, Dangl JL (2006) The plant immune system. Nature 444: 323–329

Kang JY, Nan X, Jin MS, Youn SJ, Ryu YH, Mah S, Han SH, Lee H, Paik SG, Lee JO (2009) Recognition of lipopeptide patterns by Toll-like receptor 2-Toll-like receptor 6 heterodimer. Immunity 31: 873–884

Kawasaki T, Kawai T (2014) Toll-like receptor signaling pathways. Front Immunol 5: 461

Kirschner KN, Yongye AB, Tschampel SM, Gonzalez-Outeirino J, Daniels CR, Foley BL, Woods RJ (2008) GLYCAM06: a generalizable biomolecular force field. Carbohydrates. J Comput Chem 29: 622–655

Koller B, Muller-Wiefel AS, Rupec R, Korting HC, Ruzicka T (2011) Chitin modulates innate immune responses of keratinocytes. PLoS One 6: e16594

Koymans KJ, Feitsma LJ, Brondijk TH, Aerts PC, Lukkien E, Lossl P, van Kessel KP, de Haas CJ, van Strijp JA, Huizinga EG (2015) Structural basis for inhibition of TLR2 by staphylococcal superantigen-like protein 3 (SSL3). Proc Natl Acad Sci U S A 112: 11018–11023

Krieger E, Vriend G (2015) New ways to boost molecular dynamics simulations. J Comput Chem 36: 996–1007

Lee CG, Da Silva CA, Lee JY, Hartl D, Elias JA (2008) Chitin regulation of immune responses: an old molecule with new roles. Current opinion in immunology 20: 684–689

Liu T, Liu Z, Song C, Hu Y, Han Z, She J, Fan F, Wang J, Jin C, Chang J et al (2012) Chitin-induced dimerization activates a plant immune receptor. Science 336: 1160–1164

Mack I, Hector A, Ballbach M, Kohlhaufl J, Fuchs KJ, Weber A, Mall MA, Hartl D (2015) The role of chitin, chitinases, and chitinase-like proteins in pediatric lung diseases. Mol Cell Pediatr 2: 3

Meng G, Grabiec A, Vallon M, Ebe B, Hampel S, Bessler W, Wagner H, Kirschning CJ (2003) Cellular recognition of tri-/di-palmitoylated peptides is independent from a domain encompassing the N-terminal seven leucine-rich repeat (LRR)/LRR-like motifs of TLR2. J Biol Chem 278: 39822–39829

Meng G, Rutz M, Schiemann M, Metzger J, Grabiec A, Schwandner R, Luppa PB, Ebel F, Busch DH, Bauer S et al (2004) Antagonistic antibody prevents toll-like receptor 2-driven lethal shock-like syndromes. J Clin Invest 113: 1473–1481

Morales DK, Hogan DA (2010) Candida albicans interactions with bacteria in the context of human health and disease. PLoS Pathog 6: e1000886

Morgulis S (1916) The Chemical Constitution of Chitin. Science 44: 866–867

Morris GM, Huey R, Lindstrom W, Sanner MF, Belew RK, Goodsell DS, Olson AJ (2009) AutoDock4 and AutoDockTools4: Automated docking with selective receptor flexibility. J Comput Chem 30: 2785– 2791

Netea MG, Sutmuller R, Hermann C, Van der Graaf CA, Van der Meer JW, van Krieken JH, Hartung T, Adema G, Kullberg BJ (2004) Toll-like receptor 2 suppresses immunity against Candida albicans through induction of IL-10 and regulatory T cells. J Immunol 172: 3712–3718

No HK, Cho YI, Kim HR, Meyers SP (2000) Effective deacetylation of chitin under conditions of 15 psi/121 degrees C. J Agric Food Chem 48: 2625–2627

Rosas M, Liddiard K, Kimberg M, Faro-Trindade I, McDonald JU, Williams DL, Brown GD, Taylor PR (2008) The induction of inflammation by dectin-1 in vivo is dependent on myeloid cell programming and the progression of phagocytosis. J Immunol 181: 3549–3557

Scheibner KA, Lutz MA, Boodoo S, Fenton MJ, Powell JD, Horton MR (2006) Hyaluronan fragments act as an endogenous danger signal by engaging TLR2. J Immunol 177: 1272–1281

Schmid-Burgk JL, Schmidt T, Gaidt MM, Pelka K, Latz E, Ebert TS, Hornung V (2014) OutKnocker: a web tool for rapid and simple genotyping of designer nuclease edited cell lines. Genome Res 24: 1719–1723

Stockinger LW, Eide KB, Dybvik AI, Sletta H, Varum KM, Eijsink VG, Tondervik A, Sorlie M (2015) The effect of the carbohydrate binding module on substrate degradation by the human chitotriosidase. Biochim Biophys Acta 1854: 1494–1501

To T, Stanojevic S, Moores G, Gershon AS, Bateman ED, Cruz AA, Boulet LP (2012) Global asthma prevalence in adults: findings from the cross-sectional world health survey. BMC Public Health 12: 204

Trott O, Olson AJ (2010) AutoDock Vina: improving the speed and accuracy of docking with a new scoring function, efficient optimization, and multithreading. J Comput Chem 31: 455–461

Van Dyken SJ, Liang HE, Naikawadi RP, Woodruff PG, Wolters PJ, Erle DJ, Locksley RM (2017) Spontaneous Chitin Accumulation in Airways and Age-Related Fibrotic Lung Disease. Cell 169: 497– 509 e413

Wagener J, Malireddi RK, Lenardon MD, Koberle M, Vautier S, MacCallum DM, Biedermann T, Schaller M, Netea MG, Kanneganti TD et al (2014) Fungal chitin dampens inflammation through IL-10 induction mediated by NOD2 and TLR9 activation. PLoS pathogens 10: e1004050

Wang L, Weber AN, Atilano ML, Filipe SR, Gay NJ, Ligoxygakis P (2006) Sensing of Gram-positive bacteria in Drosophila: GNBP1 is needed to process and present peptidoglycan to PGRP-SA. Embo J 25: 5005–5014

Weintz G, Olsen JV, Fruhauf K, Niedzielska M, Amit I, Jantsch J, Mages J, Frech C, Dolken L, Mann M et al (2010) The phosphoproteome of toll-like receptor-activated macrophages. Mol Syst Biol 6: 371

Woehrle T, Du W, Goetz A, Hsu HY, Joos TO, Weiss M, Bauer U, Brueckner UB, Marion Schneider E (2008) Pathogen specific cytokine release reveals an effect of TLR2 Arg753Gln during Candida sepsis in humans. Cytokine 41: 322–329

